# Scaled multidimensional assays of variant effect identify sequence-function relationships in hypertrophic cardiomyopathy

**DOI:** 10.1101/2025.05.23.655878

**Authors:** Yuta Yamamoto, Kaiser Chua, Alexis Ferrasse, Anna Kirilova, Hannah N. De Jong, Brendan J. Floyd, Christian Cadisch, Laurens Wiel, Qianru Wang, Matthew J. O’Neill, Daniel Tabet, David Staudt, John E. Goryznski, Yong Huang, Rachel H. Wilson, Arman Sharma, Althea Tapales, Rani Agrawal, Matthew T. Wheeler, Calum MacRae, Dan M. Roden, Frederick P. Roth, Andrew M. Glazer, Euan A. Ashley, Victoria N. Parikh

## Abstract

**Background:** An estimated 1 in 500 people live with hypertrophic cardiomyopathy (HCM), a disease for which genetic diagnosis can identify family members at risk, and increasingly guide therapy. Mutations in the myosin binding protein C3 (*MYBPC3*) gene account for a significant proportion of HCM cases. However, many of these variants are classified as variants of uncertain significance (VUS), complicating clinical decision-making. Scalable methods for variant interpretation in disease-specific cell types are crucial for understanding variant impact and uncovering disease mechanisms.

**Methods:** We developed a scaled multidimensional mapping strategy to evaluate the functional impact of variants across a critical domain of MYBPC3. We incorporate saturation base editing at the native *MYBPC3* locus, a long-read RNA sequencing-enabled assay of variant splice effects, and measurements of HCM-relevant phenotypes, including MYBPC3 abundance, hypertrophic signaling, and ubiquitin-proteasome function in human induced pluripotent stem cell-derived cardiomyocytes (iPSC-CMs).

**Results:** Our multidimensional mapping strategy enabled high-resolution functional analysis of *MYBPC3* variants in iPSC-CMs. Targeted transient base editing generated a comprehensive variant library at the native locus, capturing diverse variant effects on cellular HCM-relevant phenotypes. Our massively parallel splicing assay identified novel splice-disrupting variants. Integration of functional assays revealed that decreased MYBPC3 abundance is a key driver of HCM-related phenotypes. In parallel, downregulation of protein degradation was observed as a compensatory response to MYBPC3 loss of function, and novel disease mechanisms were identified for missense variants near a critical binding domain, underscoring their contribution to pathogenesis. Bayesian estimates of variant effects enable the reclassification of clinical variants.

**Conclusions:** This work provides a platform for extending genome engineering in iPSCs to multiplexed assays of variant effects across diverse disease-relevant cellular phenotypes, enhancing the understanding of variant pathogenicity and uncovering novel biological mechanisms that could inform therapeutic strategies.

## Introduction

Recently, a wealth of data informing our understanding of sequence-function relationships has emerged from multiplexed assays of variant effect^1^. The rate of production of individual variant effect estimates derived from these efforts far outpaces traditional variant testing^2^, and, when assembled into variant effect maps, uncover important aspects of protein structure and function^2,3^. They also address a critical need in precision medicine by providing evidence for or against the pathogenicity of frequently identified variants of uncertain significance (VUS, variants with limited or no evidence for disease causality that greatly diminish the diagnostic yield of clinical genetic testing)^4,5^. Proactive generation of scaled functional evidence for variant effects contributes to improved speed and accuracy of variant adjudication for individual patients^5^. In addition, integration of these functional data into increasingly sophisticated variant effect predictors will help to increase the breadth of their implementation in the clinic and laboratory^6^.

Technical challenges in high-throughput genomic mutagenesis of non-immortalized cell types (e.g., neurons and cardiomyocytes) has limited the use of scalable technologies to genes associated with tissue-specific diseases. Extralocal expression in immortalized cell lines allows for limited insights into protein abundance, but does not fully capture the biological impact of these variants. This is true, in particular, for diseases of cardiac muscle (cardiomyopathies), where maintenance of allelic stoichiometry is critical^7,8^, and where the specific biology of the myocyte drives much of the downstream mechanism of variant effect. To date, the highest resolution variant effect maps rely on immortalized non-cardiac cell lines, which limits the phenotypic assessment of variants in genes expressed specifically in cardiomyocytes. Further, the majority of these maps are generated from cDNA construct libraries rather than direct genome editing, limiting their ability to detect downstream effects of splice-altering variants or replicate genomic and cellular contexts. As precision therapies for genetic cardiovascular disease progress to first-in-human trials^9,10^, improvement in the diagnostic yield of genetic testing and knowledge of variant-specific disease mechanisms is more critical than ever.

Hypertrophic cardiomyopathy (HCM) is a prevalent monogenic cardiac disease associated with heart failure and sudden death for which genetic diagnosis is clinically actionable. Genetic testing can exclude family members from frequent and expensive clinical screening and identifies patients who will benefit from precision therapies like cardiac myosin inhibitors and gene replacement^11–13^. The majority of causative variants in HCM are identified in *MYBPC3*, which encodes myosin-binding protein C (cMyBP-C). cMyBP-C plays a critical role in the regulation of cardiomyocyte contractility, acting as a brake on the contractile activity of the sarcomere. Loss of function variants in *MYBPC3* cause hypercontractility and left ventricular hypertrophy, cardinal features of HCM^14^, however, the vast majority of missense and potentially splice-altering *MYBPC3* variants remain VUS.

To allow the depth, breadth, and throughput necessary to simultaneously evaluate hundreds of *MYBPC3* variant effects in their native genomic locus, we employed base editing and developed multiple phenotypic assays in human induced pluripotent stem cell-derived cardiomyocytes (iPSC-CMs). We combined a long-read-sequencing-based splicing assay with scaled *in situ* mutagenesis followed by multiple functional assays to simultaneously assess the effects of over 400 *MYBPC3* variants on iPSC-CM phenotypes. Integration of this multidimensional phenotyping revealed novel *MYBPC3* variant effects, uncovered the complexity of sequence-function relationships in *MYBPC3*, and achieved successful reclassification of VUS for HCM patients.

## Methods

Detailed description of materials and methods is available in the **Supplemental Material**.

## Statistical analysis

In the minigene splicing assay, the proportion of the canonical isoform was calculated for each variant to quantify splicing outcomes. Differences between each variant and the wildtype were assessed using Welch’s t-test, with false discovery rate (FDR) correction applied using the Benjamini-Hochberg (BH) method to account for multiple comparisons.

For functional assays, log₂ enrichment scores were converted to Z-scores for normalization. Group comparisons were performed using the Mann-Whitney U test, and for multiple group comparisons, statistical significance was first evaluated using the Kruskal-Wallis test. If significant (p < 0.05), pairwise Mann-Whitney U tests with Bonferroni correction were conducted to adjust for multiple comparisons. All statistical analyses were conducted in Python using the SciPy and StatsModels libraries.

## Data and code availability

The raw sequencing data from experiments have been deposited in the NCBI Sequence Read Archive (SRA) under accession number PRJNA1236899. A custom pipeline for minigene splicing assay is available on GitHub (https://github.com/AshleyLab/minigene_pipeline). The scores for all assays are available in **Table S1**. Any additional information required for reanalysis is available upon request.

## Results

### Massively parallel splicing assay identifies novel splice-disrupting variants in *MYBPC3*

To evaluate variant effects on splicing, we employed a minigene assay^15^. For these and all ensuing experiments, we focused on a region of MYBPC3 with high potential for functional disruption (The tri-helix bundle (THB), a highly conserved C-terminal region of the cardiac-isoform specific M-domain, **Figure 1A** and **1B**; **Figure S1A**) ^16,17^. We confirmed that the reference sequence minigene plasmid generated the same splice events as the endogenous *MYBPC3* gene in iPSC-CMs (**Figure S1B** through **S1D**). Next, we designed a minigene plasmid library consisting of all possible single nucleotide variants at the intron 11–exon 12 junction, each tagged with a unique barcode (**Figure S1E** and **S1F**). Using long read sequencing and a custom pipeline (see **Methods**; **Figure S1G** and **S1H**), we evaluated splicing consequences of 402 variants (93.1% of possible variants; **Figure 1C** through **1F** (iPSC-CM); **Figure S2A and S2B** (HEK293T)). To benchmark the splicing assay, we determined the proportion of reads representing canonical vs. non-canonical splice isoforms of the minigene associated with known disease causing (termed “pathogenic or likely pathogenic” (P/LP, n=7)) or unlikely to contribute to disease (termed “benign or likely benign” (B/LB, n=9)) variants (classified according to American College of Medical Genetics and Genomics (ACMG) criteria; **Table S2**)^18,19^. P/LP variants exhibited a markedly lower proportion of canonically spliced isoforms compared to B/LB variants, indicating that this assay can distinguish between disease-causing and benign *MYBPC3* variants (**Figure 1C**).

**Figure 1.**
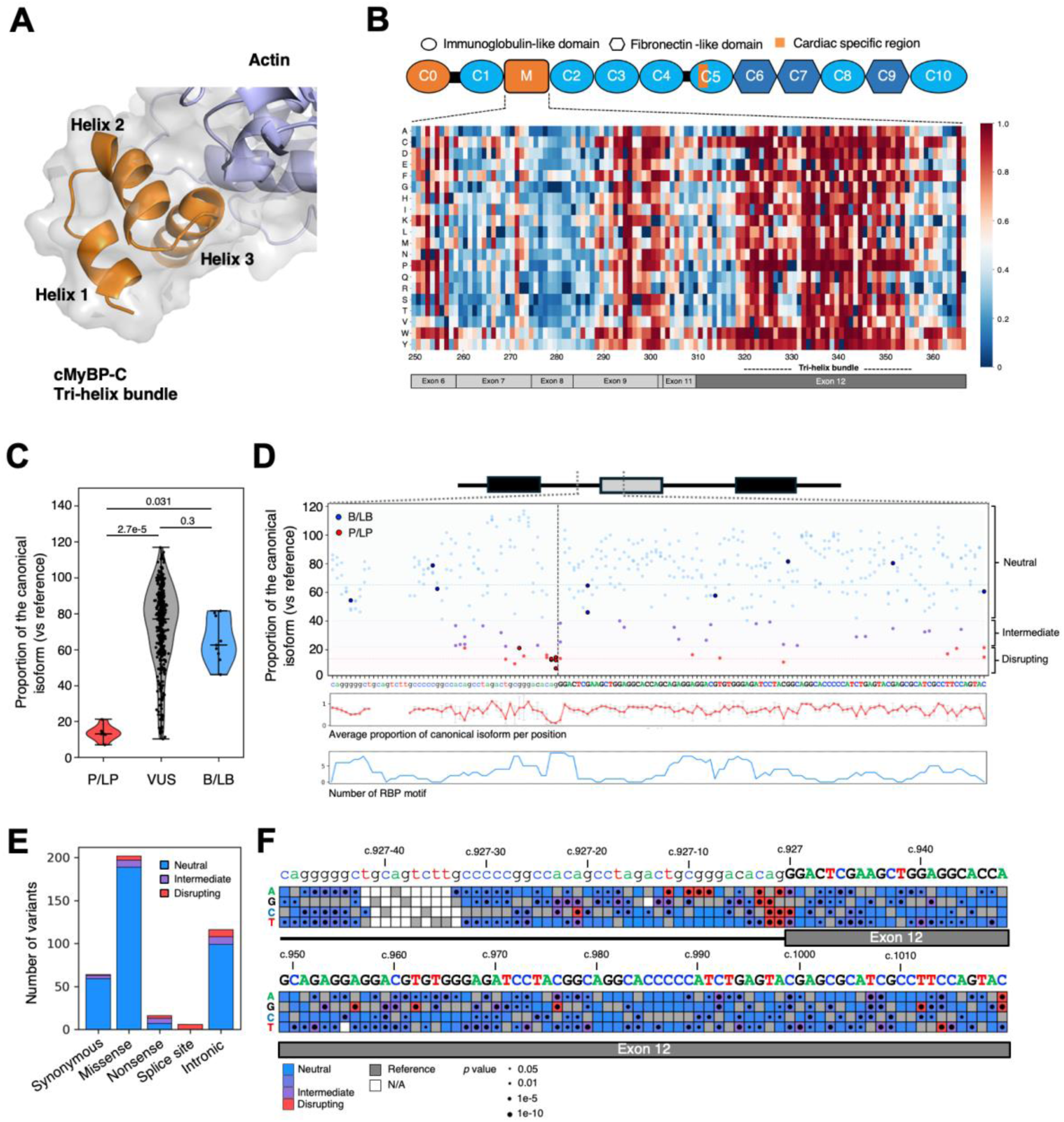
cMyBP-C M-domain and massively parallel splicing assay using minigene plasmid library. **A**, 3D structures (PDB: 7TIJ) of tri-helix bundle (THB, orange) and actin (purple). **B**, AlphaMissense prediction scores in cMyBP-C M-domain.The AlphaMissense prediction scores range from 0 (blue) to 1 (red), with higher scores indicating a greater likelihood of pathogenicity. C-terminal region of M-domain (p.319-p.356, encoded by exon 12), is a potential hotspot for pathogenic missense variants. **C**, Proportion of the canonical isoform of known pathogenic or likely pathogenic (P/LP), benign or likely benign (B/LB), and all other variants (P/LP; n = 7, B/LB; n = 9, and VUS; n = 386). Nonsense variants were excluded from classification as P/LP (see Methods). **D**, Landscape of single variant effects on canonical splicing (top). P/LP and B/LB variants are highlighted. The range of disrupting (red), intermediate (purple), and neutral (blue) are indicated by shading. Red and blue dashed lines indicate the mean proportion of canonical isoform (vs reference minigene) in P/LP and B/LB. The averaged proportion of the canonical isoform (vs reference minigene) in each position (middle). The number of RNA binding protein (RBP) motifs in each position (bottom). **E**, Number of variants classified based on proportion of the canonical isoform in each variant type. **F**, Variant effect map for splicing. X axis indicates the position in *MYBPC3* c.927-50 – c.1020. Y axis indicates nucleotide identity. Box color denotes the variant classification based on proportion of the canonical isoform. Reference nucleotides were denoted with gray squares. Blank boxes indicate missing data or variants were filtered out due to low confidence.

We classified the effects of the remaining 386 variants in the minigene library as splice-disrupting, -intermediate, or -neutral based on the 95% confidence intervals of the distributions of known splice-disrupting P/LP versus B/LB variants (see **Methods**; **Figure 2D** through **2F**; **Figure S2C**). Excluding known P/LP variants, 21 variants were classified as splice-disrupting. As expected, 6 of these 21 variants (29%) were located at canonical splice sites. In addition to these, 15 variants across both introns and exons were identified as splice-disrupting variants.

**Figure 2.**
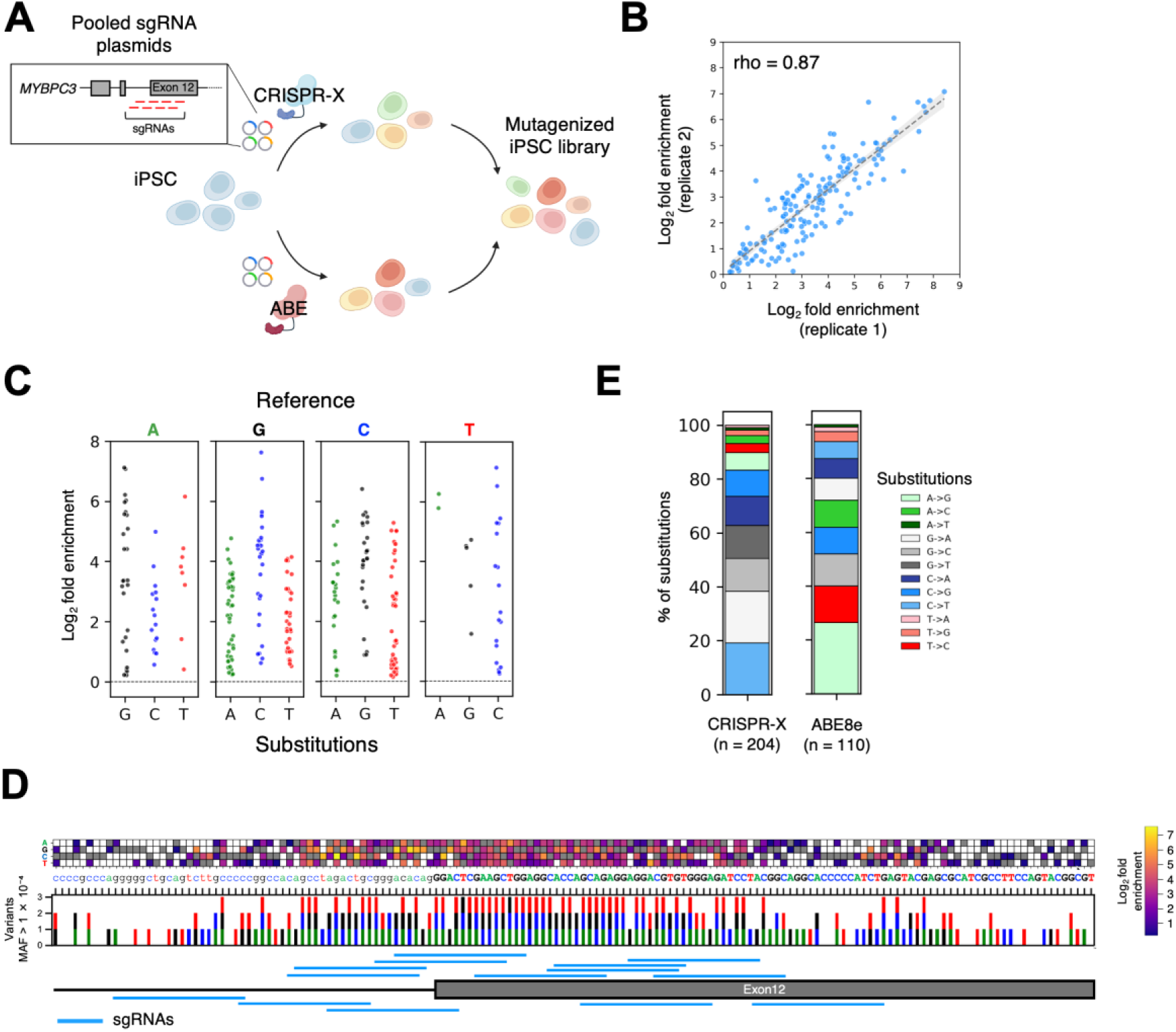
*in situ* mutagenesis using base editors. **A**, Schematic of *in situ* mutagenesis approach. CRISPR-X and Adenine base editing were performed separately. Created with BioRender.com. **B**, Enrichment scores in biological replicates (rho = 0.87, *p* = 5.14e-51). **C**, Mutagenesis efficiency of individual variants in the mutagenized iPSC library (CRISPR-X + ABE8e). Each plot represents the enrichment score for each variant. **D**, A total of 281 identified variants in mutagenized iPSC library and their enrichment scores. The color of boxes reflects the fold enrichment scores. Reference nucleotides were denoted with gray squares. Blank boxes indicate missing data or variants were filtered out due to low confidence. **E**, Proportion of nucleotide changes observed in CRISPR-X and Adenine base editor (ABE8e) library. A total of 145 and 65 variants were identified in the CRISPR-X and AEB8e libraries, respectively.

We identified 48 variants with an effect on splicing (21 splice-disrupting and 27 with an intermediate effect) in the minigene library and analyzed the frequency of each type of splicing event associated with these variants (**Figure S2D** through **S2G**). The cryptic splice sites used in those variants were also observed to be used at low frequencies in the reference sequence minigene, and this was confirmed in human cardiac tissue from a patient without an *MYBPC3* variant (**Figure S2H and S2I**). This suggests that alternative splice sites chosen by the splicing apparatus in the event of canonical site disruption are consistently recognized by the splicing apparatus, but with a lower affinity than an intact canonical splice site. As expected, SpliceAI^20^ predictions (see **Methods**) were correlated overall with variant effect (**Figure S2J**), but this correlation was much better for intronic variants than exonic variants, emphasizing the potential of functional data to inform computational variant effect predictors.

To further investigate the splicing regulation of this region, we integrated our splicing assay data with RNA-binding protein (RBP) motif data (ATtRCT, https://attract.cnic.es/documentation)^21,22^. Splice-affecting variants were enriched in positions where RBP motifs were found (**Figure 1D**). This revealed 20 RBPs potentially involved in the splicing regulation of *MYBPC3* including HNRNPA1 and SRSF5, and identified several active RBP motifs in iPSC-CMs (**Figure S2K** and **S2L**). Notably, these splicing factors have been implicated in cardiac development, hypertrophy, and heart diseases^23–26^.

### Multiplexed assays enable measurement of variant effects on HCM-relevant iPSC-CM phenotypes

To connect the direct effects of splice disruption and non-splice mediated mechanisms of *MYBPC3* variant effect to HCM-relevant phenotypes, we evaluated the effect of native locus *MYBPC3* variants on cMyBP-C abundance as well as markers of cardiomyocyte hypertrophy (brain natriuretic peptide (BNP))^17,27^ and ubiquitin-proteosome system function (UPS, impaired in *MYBPC3* HCM^28,29^) in iPSC-CMs (**Figure 3B**).

**Figure 3.**
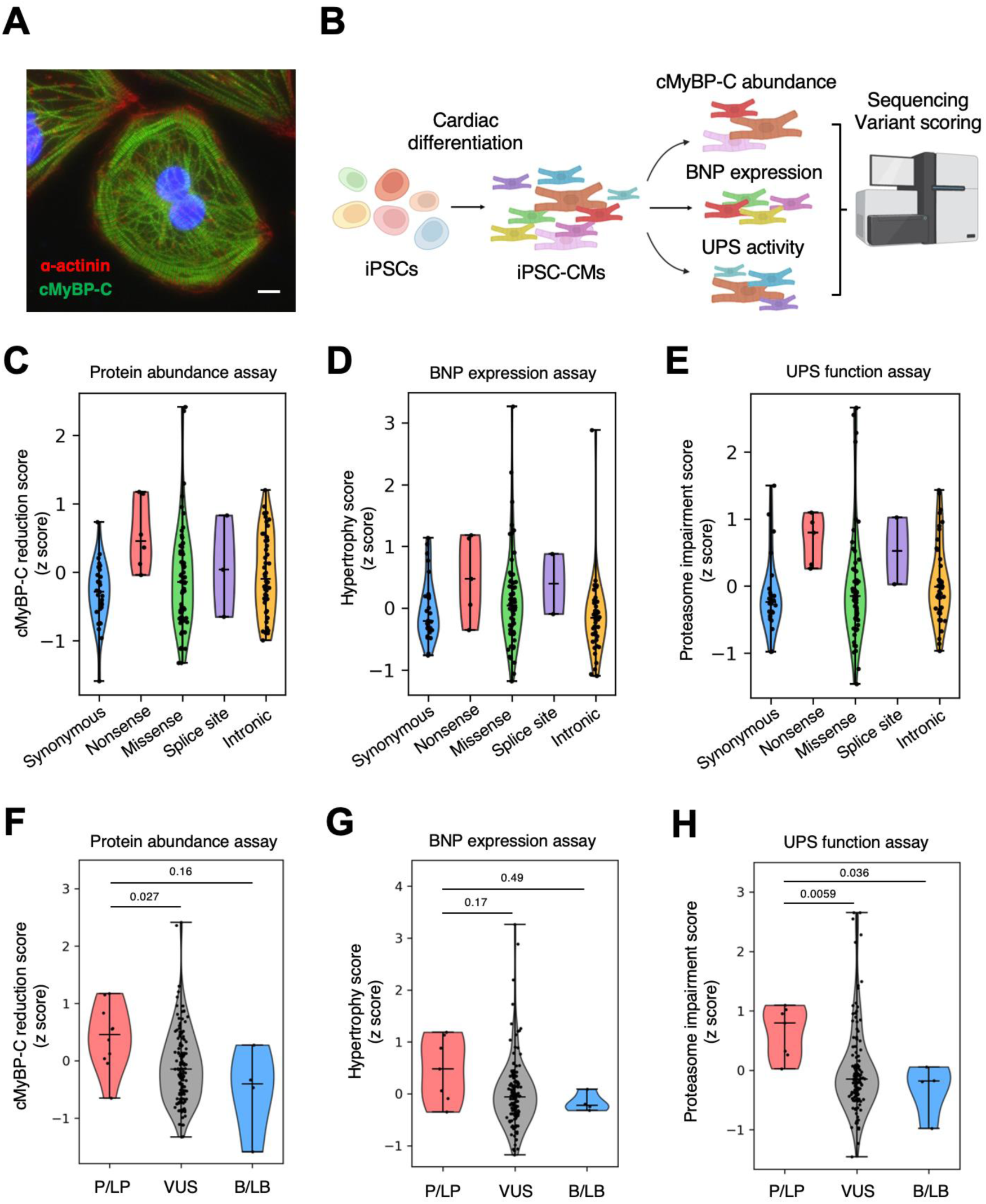
Multiplexed functional assays using iPSC-derived cardiomyocytes. **A**, Immunofluorescence image of α-actinin (red) and cMyBP-C (green) in iPSC-derived cardiomyocytes (iPSC-CMs). Scale bar: 20 µm. **B**, Schematic of selection assays design using iPSC-CMs derived from mutagenized iPSC library. Created with BioRender.com. **C** through **E**, Functional scores in each variant type. **F** through **H**, Functional scores in pathogenic or likely pathogenic (P/LP), benign or likely benign (B/LB), and all other variants (VUS).

#### Scaled base editing generates a complex variant library in iPSCs

To create a library of *MYBPC3* variants at its native locus in iPSC-CMs, we employed CRISPR-X^30^, which introduces cytidine-heavy editing, and the adenine base editor, ABE8e^31,32,33^. The pooled sgRNA plasmid library and BFP-tagged base editor plasmid were electroporated into iPSCs and Fluorescence-activated cell sorting (FACS) enriched for iPSCs with genetic modifications (**Figure 2A**; **Figure S3A**). We quantified variant frequency in each replicate compared to unedited iPSCs. Variant frequencies were highly reproducible between replicates (rho = 0.87, *p* = 1.79e-28, **Figure 2B**), and bystander edits were randomly distributed, where 0.2% of possible variant pairs were observed on the same read at a frequency higher than sequencing error (**Figure S3C** and **S3D**). In total, 281 variants were identified after stringent quality filtering (see **Methods**) by combining CRISPR-X and ABE8e mutagenesis (**Figure 2C** and **2D**), each of which introduced different genetic modifications (**Figure 2E**). These mutagenized iPSC libraries were differentiated into cardiomyocytes for multiplexed functional assays to evaluate variant effect on HCM-relevant phenotypes.

#### Multidimensional assays differentiate between known pathogenic and benign variants

To comprehensively evaluate *MYBPC3* variant effects in high-throughput, we developed FACS-based assays of (1) cMyBP-C, (2) BNP protein expression and (3) UPS function in iPSC-CMs (**Figure 3A** and **3B**) and tested these in cells harboring pathogenic *MYBPC3* mutations (**Figure S4**):

1. *cMyBP-C protein abundance (cMyBP-C reduction score)*: We measured cMyBP-C protein abundance in single iPSC-CMs by fluorescence-conjugated antibodies and FACS and compared the frequency of each *MYBPC3* variant between low- and high-cMyBP-C abundance cell populations (**Figure S5A**) Enrichment scores (cMyBP-C reduction scores) were calculated for 155 variants. As expected, nonsense variants were enriched in the low cMyBP-C population and exhibited high cMyBP-C reduction scores (**Figure 3C**). Overall, 46 variants in the region of interest had high cMyBP-C reduction scores, including 6 pathogenic variants and 4 stop-gain variants (**Figure 4A**).
2. *BNP protein expression (Hypertrophy score)*: We measured BNP expression in single iPSC-CMs by fluorescence-conjugated antibodies and FACS. The hypertrophy score of 146 variants were obtained by quantifying variant enrichment in the high-BNP cell population (**Figure 3D**; **Figure S5B**). Nonsense and splice site variants showed high hypertrophy scores, while most synonymous and intronic variants showed low scores, consistent with their presumed lack of functional impact (**Figure 3D**). Total 34 variants showed a high hypertrophy score, including 4 pathogenic variants (**Figure 4A**).
3. *UPS function (Proteosome impairment score)*: We assessed proteasome activity by transfecting cells with Ub-G76V-GFP^34^ plasmid and assessing for its clearance. Proteasome impairment scores were calculated by comparing variant frequencies between the GFP-positive (low proteasome activity) and GFP-negative (normal proteasome activity) populations (**Figure 3E**; **Figure S5C**). Nonsense and splice site variants were enriched in the GFP-positive population, indicating that these variants impaired UPS activity in cardiomyocytes. Twenty-six variants showed a high proteasome impairment score, including 4 pathogenic variants (**Figure 4A**).

**Figure 4.**
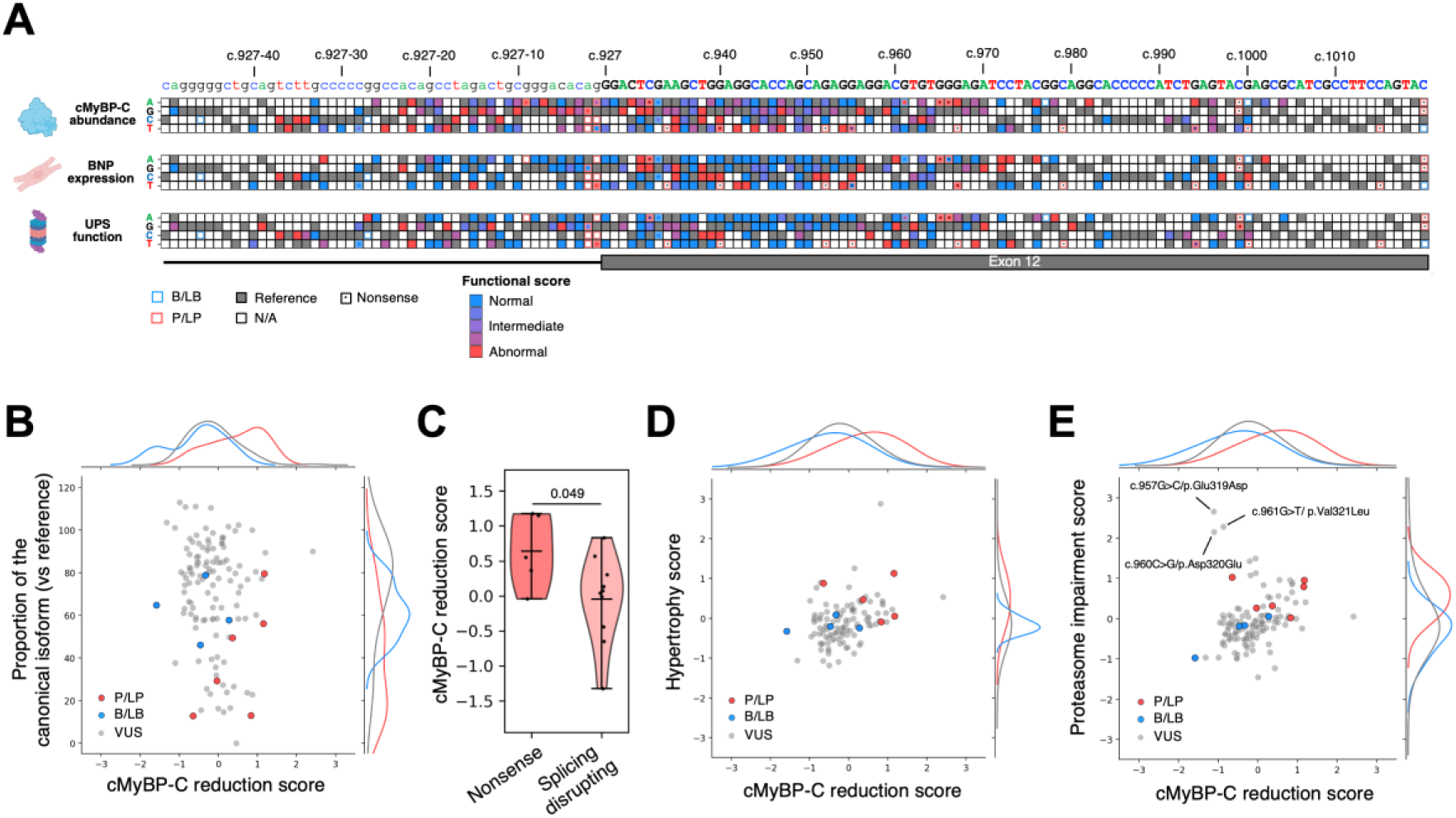
Integrated analysis of multidimensional functional assays. **A**, Variant effect map on cMyBP-C abundance (top), BNP expression (middle), and ubiquitin-proteasome system (UPS) function (bottom). The color of boxes reflects each functional score. Reference nucleotides were denoted with gray squares. Blank boxes indicate missing data or variants were filtered out due to low confidence. Nonsense variants are denoted by asterisks. Created with BioRender.com. **B,** Comparison of normalized proportion of the canonical splicing isoform and cMyBP-C reduction scores (n = 121, rho = -0.15, *p* = 0.094). **C**, cMyBP-C reduction scores in nonsense and splice-disrupting variants. **D** and **E**, Functional scores (hypertrophy and proteasome impairment) are plotted against for cMyBP-C reduction scores (BNP expression; n = 121, Proteasome impairment; n = 131).

Overall, functional assay scores aligned with anticipated profiles of different variant types. Nonsense and splice site variants exhibited high scores in each functional assay, while synonymous variants tended to show low scores (**Figure 3C** through **3E**). The wide distribution of scores for missense variants in each functional assay indicates that they represent a heterogeneous population of variants, potentially with diverse effects on each disease-relevant iPSC-CM phenotype.

To assess whether these multiplexed functional assays have sufficient sensitivity to distinguish between pathogenic and benign variants, we compared the scores of known disease causing (P/LP) and benign (B/LB) variants. Comparison of scores from each assay for P/LP variants showed a significant difference as compared to the VUS distribution in the cMyBP-C protein abundance and UPS function assays (*p*<0.05 for both assays, **Figure 3F** through **3H**). The log-likelihood ratio of disease causing (P/LP) and benign (B/LB) variants in each assay was used to classify variant effects as normal, intermediate, or abnormal in each assay (**Figure S5D** through **S5I**).

### Multidimensional assays of variant effect identify multiple underlying disease mechanisms

To understand the mechanism by which individual *MYBPC3* variants might contribute to HCM, we compared the results of each functional assay (**Figure 4A**; **Figure S6A** and **S6B**). There was not a clear correlation between cMyBP-C protein reduction and splice disruption scores (rho = -0.15, *p* = 0.094, **Figure 4B**; **Figure S6C**). Nonsense variants *did* display overall more cMyBP-C reduction than splice disrupting variants (**Figure 4C**). Supporting the effect of reduced cMyBP-C abundance on cardiomyocyte hypertrophy signaling, there is a modest correlation between hypertrophy score and cMyBP-C reduction score for each variant (**Figure 4D**, rho = 0.37, *p* = 3.51e-5). We also found a *positive* correlation between proteasome impairment and cMyBP-C reduction scores (**Figure 4E**, rho = 0.38, *p* = 7.38e-6) consistent with prior reports in single variants^35^. To interrogate whether this correlation reflects the functional deactivation of UPS in response to imbalanced allelic stoichiometry, we regressed the proteasome impairment score against the proportion of the canonical isoform identified in association with each variant. We found only a weak correlation between the proportion of the canonical isoform and proteasome impairment score (rho=-0.19, *p* = 0.035, **Figure S6D**), indicating that proteasome impairment is driven by additional disease mechanisms independent of mis-splicing and nonsense mediated decay in *MYBPC3* HCM.

Our region of interest contains a highly conserved and structured region known as the tri-helix bundle (THB, p.319 - 356), which may serve as a potential interaction platform for actin and calmodulin^36,37^. The mean functional scores of single amino acid substitutions were mapped onto the one-dimensional structure of cMyBP-C (**Figure 5A**, 13 splice-disrupting variants excluded). We compared functional scores of variants within the THB region to those outside it (**Figure 5B** through **5D**). We found that variants located within the THB region impair proteasome activity more extensively than those outside the THB region (**Figure 5D** and **5E**). Although a similar trend was observed for hypertrophy scores, the difference did not reach statistical significance (**Figure 5C**). In contrast, the impact of missense variants on protein abundance levels was not found to be associated with specific structural regions (**Figure 5B**). Several missense variants in helix 1 of the THB (c.957G>C/p.Glu319Asp, c.960C>G/p.Asp320Glu, and c.961G>T/p.Val321Leu) showed intermediate or low splicing and cMyBP-C reduction scores, but high proteasome impairment scores (**Figure 4E**). To investigate the potential mechanism of these missense variants, we used FoldX v5.1^38,39^ to predict the ΔΔG of these variants vs reference sequence. In the majority of the NMR states, the missense variants are not thermodynamically favored, indicating a likelihood of protein instability which may act as a loss of function mechanism (**Figure S7**). We also found that these missense variants in helix 1 of the THB domain caused conformational changes that may disrupt cMyBP-C binding to actin or calmodulin, or altering calcium sensitivity, as demonstrated for a missense variant at residue 322 in a previous study^40^(**Figure 5F** through **5I**).

**Figure 5.**
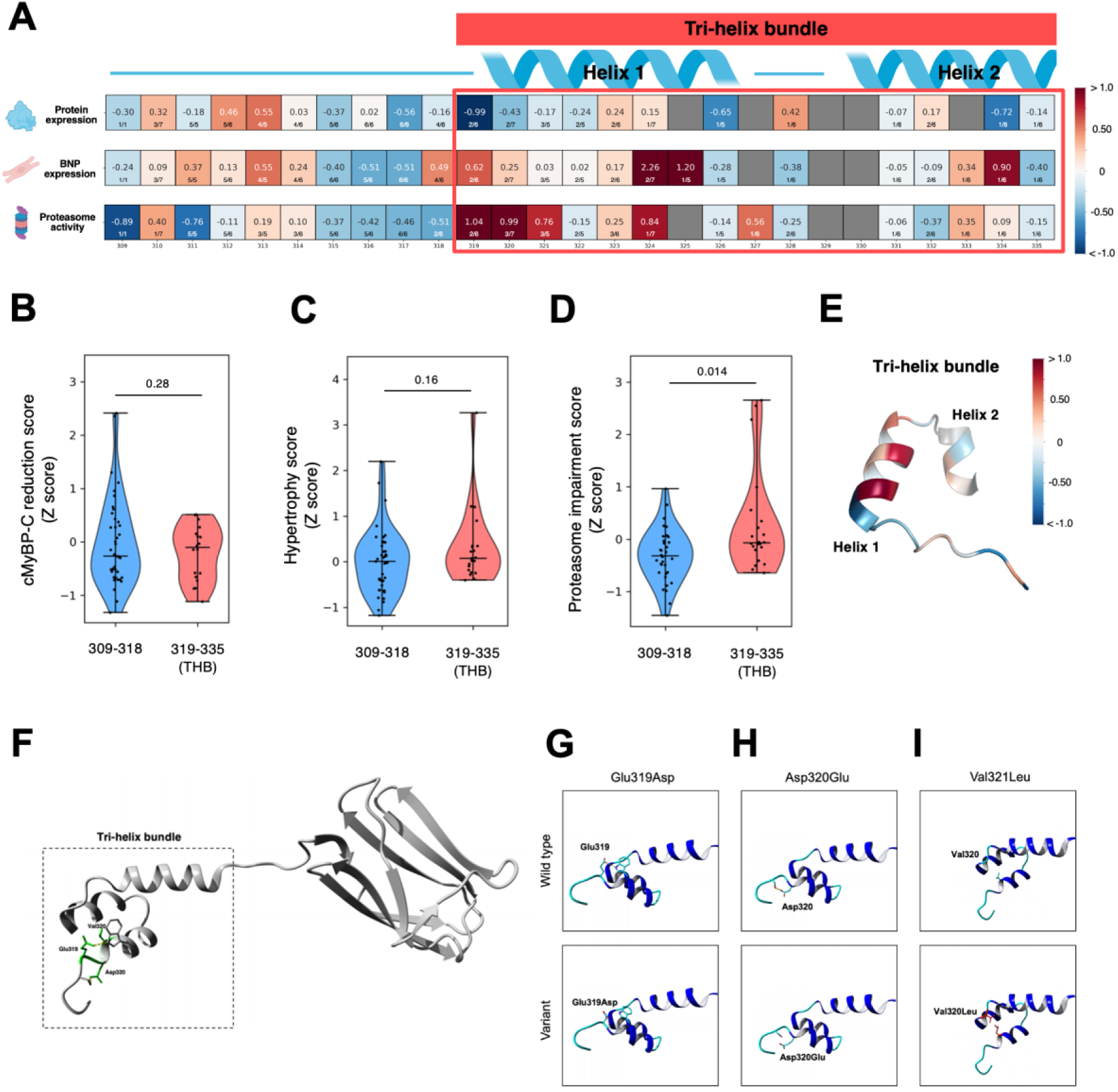
Multidimensional assay decodes missense variants of *MYBPC3*. **A**, Mean functional scores of single amino acid substitutions in 1D structure of cMyBP-C. Each cell represents an amino acid position, with the background color and numerical value indicating the mean functional score. The fraction below (e.g., 2/6) represents the coverage of observed substitutions (number of the substitution was observed / total possible substitution for that position). Gray indicates missing data or variants were filtered out due to low confidence. **B** through **D**, Comparison of functional scores between variants within and outside of the tri-helix bundle (THB) region. **E**, Mean proteasome impairment score of amino acid substitutions were utilized to map onto 3D structure of the THB. **F**, Structure of cMyBP-C M-domain. **G** through **I**, Local atomic interaction in wild type and mutated structures for Glu319Asp, Asp320Glu, and Val321Leu. The p.Glu319Asp variant disrupts the salt bridge between p.Glu319 and p.Trp322, and creates a new salt bridge with the structure’s backbone at position 319. The p.Asp320Glu variant causes a loss of the salt bridge formed with p.Ser318. The missense variant p.Val321Leu introduces a much larger residue at position 321 which overlaps with the molecular surface of p.Val342 and thus will not fit without a conformational change.

### Multidimensional assays of variant effect provide clinical insight

To assess the strength of evidence for variant interpretation provided by each of our assays, we calculated the odds of pathogenicity scores for all variants observed at high frequency in the assay (OddsPath, a recommendation-based Bayesian estimate of assay evidence strength based on the scores of known P/LP and B/LB variants, **Tables S2** and **S3**)^41^. We incorporated these data into clinical interpretation of 17 variants identified in the Sarcomeric Human Cardiomyopathy Registry of HCM (SHaRe, containing high resolution clinical data on 17,000 patients ^42,43^, **Table S4**) (**Figure 6A and 6B**). For known pathogenic and likely pathogenic (P/LP) variants (n=8), ACMG classifications based on prior data were unchanged after addition of assay data. There were 8 variants classified as VUS prior to application of assay findings. Contribution of functional data from the assays reclassified 2 of these to likely benign, and lowered suspicion for pathogenicity in an additional 4. One of these VUSs, c.1008C>T (p.Ile336Ile) was of particular interest as it had just been reported in a pediatric case of severe HCM^44^. Supporting evidence from the splicing assay significantly raised suspicion for its pathogenicity (alongside its pathogenic neighbor, c.1007T>A (p.Ile336Asn, **Figure 6C**)). As these assays are extended to include additional pathogenic and benign variants, the strength of evidence of these predictions is likely to increase, making them even more impactful in variant interpretation.

**Figure 6.**
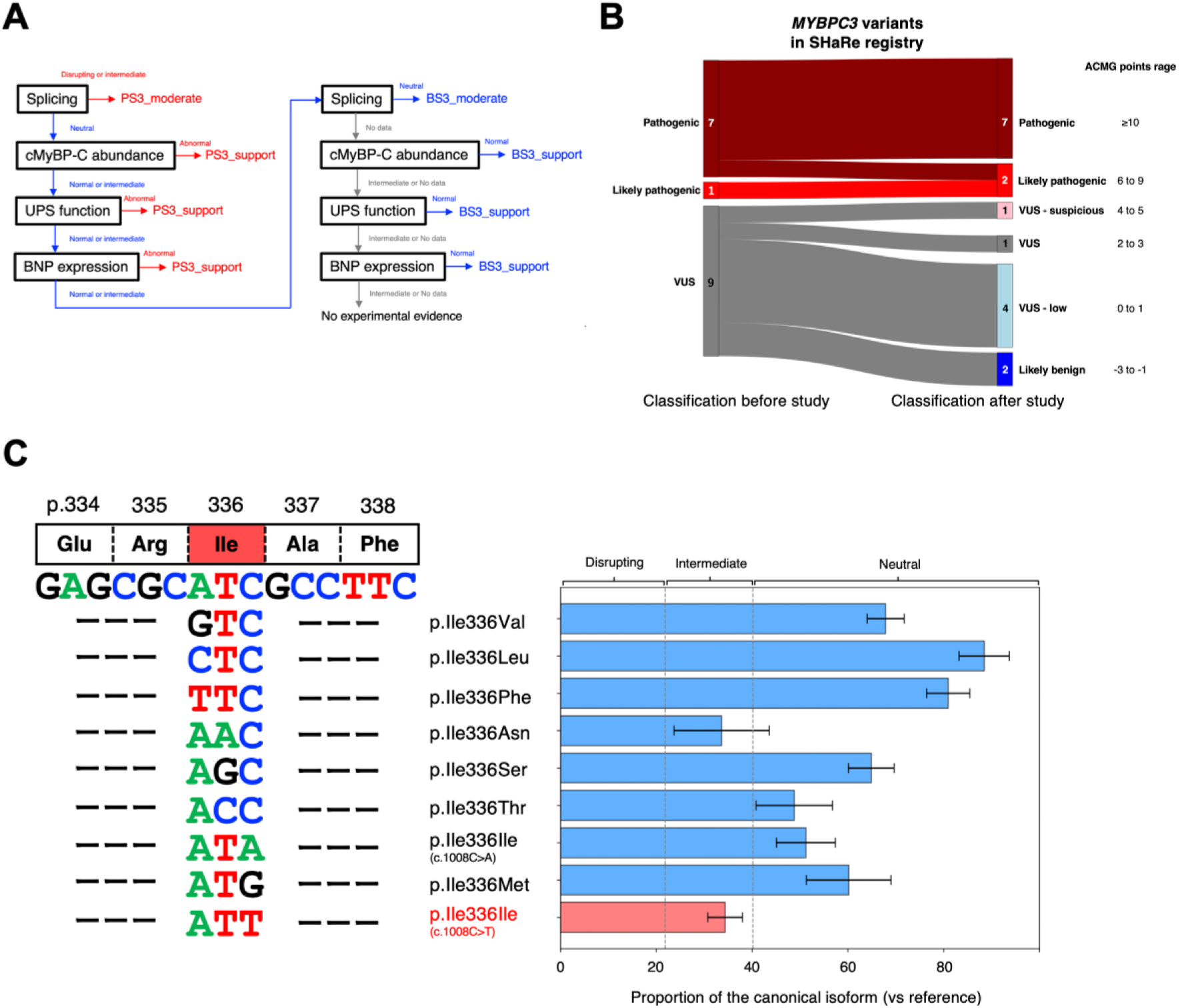
Functional assay evidence-based variant classification. **A,** Decision tree for the variant classification for clinical interpretation using multidimensional functional assay data. We integrated data across assays by determining the assay with the strongest evidence for pathogenicity, and, failing that, the strongest evidence for benign classification as has been suggested for incorporation of Bayesian variant effects into clinical variant interpretation. The strength of evidence in each assay is based on Odds of pathogenicity (OddsPath). **B,** Classification of 17 variants (7 pathogenic, 1 likely pathogenic, 9 variants of uncertain significance (VUS)) identified in the Sarcomeric Human Cardiomyopathy Registry (SHaRe) based on our functional assay data. These variants were adjudicated by two independent cardiovascular genetics specialists with and without the functional data provided by our assays according to ACMG guidelines using a validated points system for reproducibility. To combine the data from our four assays, we assessed evidence for pathogenicity first. In the case of conflicting evidence, given the different disease mechanisms presented, the strongest level of evidence available was accepted. If none of the assays showed evidence of pathogenicity, the strongest available evidence for benign effect was accepted. **C,** Single-nucleotide resolution assay revealed the splicing disruption in the synonymous variant (c.1008C>T/lle336lle) identified in SHaRe registry.

## Discussion

We used targeted base editing and long-read sequencing in induced pluripotent stem cell derived cardiomyocytes to extend a tissue-specific, disease-relevant model to multidimensional assays of variant effect within the native genomic context. We achieved multidimensional interrogation of over 400 intronic and exonic variants in the THB region and the intron11-exon12 junction of *MYBPC3.* Large-scale combinational functional assays were sensitive enough to distinguish between known benign and pathogenic variants identified in HCM patients, and to classify VUS. We show the diversity and heterogeneity of individual variant effects on HCM-relevant phenotypes. These underscore that a reduction of protein abundance is a major disease mechanism, closely tied to assays of cellular hypertrophic signaling and UPS function, but that UPS function can be impaired by some missense variants even in the absence of abnormal splicing or reduced cMyBP-C abundance. The variant effects identified here are immediately translatable to the clinic where they can help to resolve VUS for HCM patients.

The variant library construction and multi-assay application methods we present create an opportunity for high-depth phenotyping of variants in cardiomyocyte-specific genes at the genomic locus in a cardiomyocyte context. Prior variant effect measurements in a cardiomyocyte context have made significant progress in studying multiple variants simultaneously either in mid-throughput series^45^, or by introduction of landing pads and gene silencing of the specific region of interest in the model line^46,47^. The method presented here can be applied at any genomic locus of interest, and can be studied in pooled assays with broadly disease-relevant phenotypes such as UPS function and BNP expression. iPSCs, with their ability to differentiate into various cell types, hold promise across many tissue-specific genetic diseases. In the future, the rapid adoption of prime editing and other high efficiency methods of editing in iPSC contexts presents an opportunity to characterize large-scale variant libraries introduced at the native locus^47–50^. Our study advances splice reporter assays^15^ by combining long-read sequencing with a novel analysis pipeline: cross-platform subassembly, which extends prior algorithms for barcoded variant libraries^15,51^, and quantifies full-length isoforms by VIsoQLR^52^ and link variants to alternative splicing events. This approach not only extends splice-effect mapping deeper into exons, but also provides novel biological insights in splicing regulation.

Integration of multiple functional assays using iPSC-CMs provided novel insights into both the interpretation of variant pathogenicity and the sequence-function mechanisms. While haploinsufficiency remains the primary pathogenic mechanism for *MYBPC3* variants, consistent with previous studies using iPSC-CMs^17,35^, we observed splice-disrupting variants that maintained normal-intermediate protein abundance levels, rendering them undetectable as high-effect variants when assessed solely via cMyBP-C protein abundance. This observation underscores the importance of complementary functional assays to provide a more comprehensive understanding of variant effects. Prior work has posited that decreased protein clearance via reduced UPS function reflects an innate capacity to maintain normal cMyBP-C levels through a compensatory slowing of cMyBP-C degradation^35^. In concordance with this, we found a correlation between protein reduction scores and proteasome impairment scores, in particular for nonsense variants. UPS dysfunction may be induced by multiple factors, including an increased accumulation of misfolded proteins, an elevated demand for protein turnover due to functional abnormalities such as hypercontraction, and compensatory mechanisms to address haploinsufficiency. Of note, this partial maintenance of protein levels was more pronounced in splicing variants compared to nonsense variants, indicating the possibility that protein levels are partially maintained through the transcription of a small fraction of canonical isoforms from the variant allele at the upstream mRNA expression level. At the same time, we uncover missense variants in the THB that may cause MYBPC3 loss of function *without* altered splicing or reduced protein abundance and identify two potential underlying mechanisms using prediction of changes in structural free energy and three dimensional conformation changes. These observations indicate that, while protein abundance levels alone are insufficient to fully characterize variant pathogenicity, the integration of protein abundance, splice readouts, and UPS function can unlock more variant effects and new mechanisms of disease.

The findings of our study exist in the context of its necessary limitations. While this study provides a high-depth characterization of over 400 variants in a critical gene causing HCM, it is limited to the M-domain. Extension of this work to the entirety of *MYBPC3*, as with any large gene, will require sizable investment, though the technologies presented here are intended to reduce this significantly. Wide-window base editing carries the risk of off target or bystander edits. We have reduced the risk of this significantly through transient expression of editors and find a very low frequency distribution of paired bystander edits. Validation against *a priori* knowledge of the several pathogenic variants in this region shows that the signal to noise ratio of our assays is high. Notably the base-pair resolution achieved by this method adds additional precision to previously reported base editing methods in which intended editing windows rather than variants were associated with assay outcome^53^. The rigor required for application of variant effect assays to variant interpretation is replicated here in our OddsPath analysis, and the ratio of high effect to low effect VUS we identify is expectedly skewed toward low effect-size variants.

The multidimensional evaluation of hundreds of individual variants here demonstrates the complexity underlying disease mechanisms and the need for a nuanced approach to functional evaluation, even within a single domain of a single disease gene. We have demonstrated that transient expression of base editors in human induced pluripotent stem cells is a feasible and broadly applicable technology for studying variant effects in cardiomyopathy, and immediately extendable to other derived primary cell types. Our maps identify exonic variants that affect splicing and are not currently identified by deep learning enabled prediction algorithms. Combining multiple assays with variant-specific protein structure modeling, we identify variants with high effect that may act through a different mechanism than previously described for this gene. Our findings are immediately applicable in HCM patients. This work demonstrates the breadth of impact of disease-relevant variant effect mapping and extends current technology to previously intractable models of disease.

## Acknowledgements

We thank the Ashley laboratory members for critical discussion and feedback. We are grateful to Dr. Micael Bassik (Stanford University) for providing CRISPR-X plasmid for this project. Cell sorting/flow cytometry analysis for this project was done on instruments in the Stanford Shared FACS Facility (RRID: SCR_017788). We acknowledge the SHaRe registry for providing a list of *MYBPC3* variants from their highly clinically phenotyped registry as previously reported^43^.

## EXPANDED METHODS

### Cell lines

hiPSC lines were obtained from the Stanford Cardiovascular Institute Biobank. Patients were enrolled under the Stanford Institutional Review Board and StemCell Research Oversight Committee guidelines. All volunteers gave consent.

### Cell culture and cardiac differentiation

HEK293T cells were cultured in Dulbecco’s Modified Eagle Medium (DMEM) supplemented with 10% fetal bovine serum (FBS) at 37°C in a humidified atmosphere with 5% CO₂.

Two iPSC lines derived from healthy individuals and three iPSC lines derived from patients with HCM carrying a *MYBPC3* mutation were obtained from the Stanford Cardiovascular Institute Biobank. These iPSCs were maintained at 37°C in 5% CO₂ using mTeSR medium (STEMCELL Technologies). iPSCs were differentiated into cardiomyocytes following previously described methods^54^.

On days 9 to 11, iPSC-CMs were maintained in a lactate culture medium to enrich for cardiomyocytes. On day 12, cells were dissociated using Accutase (Sigma-Aldrich) and TrypLE Select Enzyme 10X (Thermo Fisher Scientific) and replated onto Matrigel-coated (Corning) 12-well plates. The culture medium was changed every other day until the cells were ready for functional assays at 40 days post-differentiation.

### Human heart tissue

Human transplant heart sample are collected by tissue biobank under IRB#32769.We get 1X1cm of left ventricle and the tissue is sliced using a microtome. The tissue slices were used for RNA extraction immediately after acquisition.

### Variant effect predictors SpliceAI

We compared the splicing scores (normalized_canonical_isoform) obtained from our minigene assay with the prediction scores generated by the deep learning algorithm SpliceAI^20^. SpliceAI was run with default parameters using the GENCODE annotation^55^ for the human genome version hg38 as the reference. The SpliceAI prediction score was calculated using four prediction scores—AG (acceptor gain), AL (acceptor loss), DG (donor gain), and DL (donor loss)—following the formula: SpliceAI prediction score = 1-((1-(AG)) x (1-(AL)) x (1-(DG)) x (1-(DL))

### AlphaMissense

AlphaMissense predictions for all single amino acid substitutions in the human proteome data were downloaded from Google Cloud via the link provided in the official AlphaMissense GitHub repository, decompressed, and searched using the Vim text editor for the MYBPC3 accession number/UniProt ID Q14896. The MYBPC3 predictions were extracted into a separate file for further analysis.

### SHaRe registry data

We use the Sarcomeric Human Cardiomyopathy Registry (SHaRe) to obtain *MYBPC3* variants identified patients with HCM^42^. The information of variants was exported from October 2024.

### Variant classification framework for *MYBPC3*

Variants were classified according to ACMG criteria with quantification of pathogenicity as per Tavtigian et al, 2020^56^. The specific criteria were adjusted based on the ClinGen Variant Curation Expert Panel guidelines for hypertrophic cardiomyopathy^57^ with further adjustments made for *MYBPC3* as follows. BS1 was applied for gnomAD v4 MAF > 0.0002. PM2 was applied if the variant was absent in gnomAD v4 or if the MAF were <0.00004^57^. BP4 or PP3 were applied utilizing REVEL and CardioBoost scores for missense variants (PP3_Strong if REVEL score > 0.932; PP3_Moderate if 0.773 < REVEL score < 0.932; PP3_Supporting if 0.644 < REVEL score 0.773 or CardioBoost_CM score > 0.9; BP4_Supporting if REVEL score < 0.15 or CardioBoost_CM score < 0.1)^58,59^. SpliceAI was applied for variants impacting splicing (PP3_Supporting if SpliceAI score > 0.5)^20^. PVS1 was applied for nonsense variants and canonical splice site variants within 2 nt of the exon-intron junction, as truncating variants are believed to lead to MYBPC3 haploinsufficiency and are therefore likely to be pathogenic^35^. PS3 was applied based on Ito et al^60^ with modification to PS3_Supporting based on Brnich et al^41^. PS4, PS4_Moderate, or PS4_Supporting were applied when the count of carriers in the SHaRe HCM database was at least 15, 6, or 2 individuals, respectively. The variant classification was determined using the sum of the variant scores as per Tavtigian et al 2020^56^. For the benchmarking analysis of the splicing assay, nonsense variants were excluded from classification as P/LP, as their haploinsufficiency is thought to be due to early termination of translation rather than splice disruption.

### Minigene plasmid library

We generated a reference sequence (wild type) *MYBPC3* exon 12 (164bp) minigene plasmid based on the pAG424 plasmid, following the approach described in a previous study^15^. This minigene plasmid contains *MYBPC3* exon 12 (164bp) along with approximately 200 nucleotides (nt) of flanking intronic regions (**Figure S1C**).

Based on the reference sequence, sequences encompassing all possible variants at the intron 11–exon 12 junction (from c.927-50 to c.1020, *n* = 432) were synthesized, cloned into pAG424, and tagged with a 12-nt barcode by Twist Biosciences (South San Francisco, CA, **Figure S1E**). A total of 5 ng of the minigene library DNA was used to transform E. coli DH5α competent cells (Invitrogen). Diluted replating was performed to ensure the desired barcode diversity (∼10 barcodes per variant) within the minigene library. The pooled plasmid DNA was then isolated as the minigene library using a Maxiprep kit (MACHERY-NAGEL).

### Long read DNA-seq for variant subassembly

Using nanopore long-read sequencing, we linked variants and barcodes information. The primers used for amplification are provided in **Table S5**. We amplified the sequences containing variants and barcodes from the minigene library via PCR using PrimeSTAR MAX DNA Polymerase (Takara). PCR was performed under the following conditions: 20 cycles of 98°C for 10 s, 55°C for 15 s, and 72°C for 1 min. Following amplification, the PCR amplicons were extracted from a 1% agarose gel using the QIAquick Gel Extraction Kit (QIAGEN). The purified PCR product was quantified using the Qubit dsDNA HS Assay Kit (Thermo Scientific), and a total of 200 fmol of the PCR product was used for library preparation following the manufacturer’s SQK-LSK114 protocol. The final sequencing library was loaded onto a R10.4.1 flow cell and sequenced using the PromethION platform (Oxford Nanopore Technologies). Raw sequencing data was basecalled using Dorado SUP.

### Transfection with minigene library

For HEK293T cell transfections, HEK293T cells were seeded at 40–50% confluency in a 6-well plate on the day of transfection. The minigene plasmid library was transfected into about 3 million HEK293T cells using Lipofectamine 3000 (Thermo Fisher Scientific), following the manufacturer’s protocol.

For iPSC-CM transfections, around 3-5 million iPSC-CMs were dissociated using Accutase and TrypLE Select Enzyme 10X and replated onto a 12-well plate three days prior to transfection^61^. The minigene plasmid library was transfected into iPSC-CMs using ViaFect (Promega).

### RNA extraction and library preparation for long-read RNA-seq

RNA was extracted from cells 24 hours after transfection using the RNeasy Plus Mini Kit (QIAGEN). cDNA was synthesized from 200 ng of RNA per sample using SuperScript IV (Invitrogen), with a *MYBPC3* minigene plasmid-specific primer for reverse transcription. cDNA samples were indexed by PCR and purified using the QIAquick Gel Extraction Kit. The purified cDNA was quantified using the Qubit dsDNA HS Assay Kit, and a total of 200 fmol of the PCR product was used for library preparation as described in Long read DNA-seq.

## Computational pipeline

### Sub-assembly (analysis of minigene DNA-seq data)

The full computational pipeline is diagrammed in **Figure S1G**. Quality control was performed on the FASTQ file was processed using Chopper, retaining only reads with an average base quality score of q > 10. Each randomly generated 12-nt barcode was flanked by fixed 8-nt prefix and 8-nt suffix sequences, forming a 28-nt sequence with a central 12-nt variable region. Using the reverse complement of this 28-nt sequence, reverse-complemented reads in the FASTQ file were identified using Python’s re (regular expression) module, and then converted into forward reads by reverse complementing the sequence and inverting its base quality string.

After quality control, a total of 17 million reads were retained for downstream analysis. The exact 12-nt sequences of all barcodes were identified using the forward 28-nt sequence, and the barcodes were ranked in descending order based on their frequency (i.e., the number of reads in which they appeared). The barcode’s rank was plotted against its frequency in a barcode rank plot (**Figure S1H**), where the knee point of the curve is automatically computed, serving as a threshold to filter out low-fidelity barcodes based on rank. The remaining high-fidelity barcodes were compiled into a list and used to demultiplex the reads in the input FASTQ file into new {barcode}_FASTQ files. Each {barcode}_FASTQ file was aligned to the reference amplicon sequence using minimap2, generating SAM files, which were then converted to BAM files using samtools. DeepVariant^62,63^ was used to perform variant calling on each {barcode}_BAM file, outputting a VCF file containing the variants identified for each unique barcode. Only single-nucleotide variants (SNVs) located within the mutagenesis region and marked as “PASS” in the FILTER field were retained from the VCF files. Non-unique barcodes with more than one SNV remaining after VCF filtering were excluded from further analyses. The remaining barcode. VCF files were merged and grouped by SNVs, with unique barcodes aggregated into a list and their frequencies summed for each SNV.

### Post-assembly (analysis of RNA-seq data)

Quality control using Chopper was similarly performed on the input FASTQ file, and approximately 0.5 million reads per sample were retained for downstream analysis. Thereafter, {sample}_FASTQ files were generated by demultiplexing the reads in FASTQ file using the unique index sequences of each sample replicate. A second round of demultiplexing was then conducted on each {sample}_FASTQ file using the list of all unique barcodes from the sub-assembly phase. This process produced {sample}_{barcode}_FASTQ files, which were subsequently sorted into directories named according to the SNV associated with each barcode. GMAP was used to align each {sample}_{barcode}_FASTQ file, generating corresponding {sample}_{barcode}_gff3 files.

To identify isoforms generated for each variant per sample, VIso-QLR was translated into Python and optimized for enhanced performance and scalability within the pipeline^52^. For downstream analysis, we removed isoform with < 1% proportion and variant with < 150 reads in total because of low confidence. To evaluate variant effect on canonical splicing event, we calculated the normalized proportion of the canonical isoform as follows:

Proportion of the canonical isoform = canonical isoforms/total isoforms

Normalized proportion of the canonical isoform = Proportion of the canonical isoform in variant/Proportion of the canonical isoform in reference minigene

Finally, visualization plots of isoforms and splice site usage across the mutagenesis region were generated using Matplotlib, Seaborn, and Plotly.

### RNA binding protein site prediction

To identify potential RNA-binding protein (RBP) binding sites along the *MYBPC3* reference sequence, we used the ATtRACT database (https://attract.cnic.es/documentation)^21^. Duplicate binding sites reported by different experimental approaches were removed. Binding sites overlapping by more than 2-nt for each RBP were merged. To identify only active RBPs in this region, we retained RBPs with binding sites in which more than 50% of positions were associated with splicing-affecting variants (**Figure S2K and S2L**).

### In situ mutagenesis by base editing

To introduce variants into the target region of *MYBPC3*, we designed 14 single-guide RNAs (sgRNAs) using the web-based tool CHOPCHOP (https://chopchop.cbu.uib.no/) (**Table S6**). The sgRNAs were cloned into our custom pBlueScript vector encoding the U6 promoter and gRNA scaffold via Golden Gate cloning. The cloning products were transformed into E. coli DH5α competent cells.

After verification by Sanger sequencing, each gRNA plasmid was purified using a Midiprep kit (MACHERY-NAGEL).

Just before electroporation, more than 10 million iPSCs were dissociated using Accutase following a 1-hour treatment with CloneR (STEMCELL Technologies). Base editor plasmids, CRISPR-X and ABE8e (Addgene, #138495), were engineered to include BFP as a reporter linked via an IRES sequence. These modified base editor plasmids, along with the pooled gRNA plasmid library, were electroporated into dissociated iPSCs (∼1 million cells resuspended in 100 µL of resuspension buffer, using 7.5 µg of base editor and 7.5 µg of the pooled gRNA plasmid library per zap).

After 24 hours, iPSCs were dissociated and sorted by FACS to select BFP+ cells. Five biological replicates were generated for each base editor separately. The final mutagenized iPSC library was prepared by combining the CRISPR-X and ABE8e libraries.

## High-throughput functional assays

### Immunostaining for cMyBP-C protein abundance and BNP expression assay

The samples for BNP staining were treated with Brefeldin A for 4 hours at 37°C before dissociation. About 10 million iPSC-CMs were dissociated using Accutase and TrypLE with DNase. After neutralization with culture medium, single cells were fixed with 4% paraformaldehyde for 15 minutes at room temperature and permeabilized with 0.1% saponin. The samples were stained with the following primary antibodies:

- Mouse anti-α-actinin (1:1000, Sigma) and rat anti-proBNP (1:500, R&D), or
- Rabbit anti-α-actinin (1:100, Abcam) and mouse anti-MYBPC3 (1:50, Santa Cruz)

The secondary antibodies used were:

- Goat anti-mouse Alexa Fluor 488 (1:1000, Thermo Fisher Scientific) and goat anti-rat Alexa Fluor 647, or
- Goat anti-rabbit Alexa Fluor 488 and goat anti-mouse Alexa Fluor 647. Stained iPSC-CMs were resuspended in 100–300 µL of FACS buffer (HBSS buffer with EDTA) for sorting.

### Ub-GFP plasmid transfection for UPS function assay

Using the methods described above, Ub-G76V-GFP (Addgene) was transfected into about 10 million iPSC-CMs. Forty-eight hours after transfection, the iPSC-CMs were dissociated and resuspended in 100–300 µL of FACS buffer for sorting.

### Fluorescence-activated cell sorting

Cells were sorted using a FACSAria II Cell Sorter (BD) at the Stanford Shared FACS Facility. For gating, unstained samples or non-transfected control samples were used as negative controls to define fluorescence thresholds. Sorting was performed based on specific strategies tailored to each assay to isolate the target cell populations (**Figure S5A** through **S5C**). Following sorting, the cells were resuspended in culture medium supplemented with 1% penicillin-streptomycin for continued culture or frozen at −80°C for DNA extraction.

### Extraction of genomic DNA

Genomic DNA (gDNA) was extracted from at least 4 million iPSCs using the QIAamp DNA Blood Mini Kit (QIAGEN) and from all sorted iPSC-CMs using the Puregene DNA Isolation Kit (QIAGEN). The extracted gDNA from iPSCs was quantified using the Qubit BR Kit (Thermo Fisher Scientific), while the gDNA from sorted iPSC-CMs was quantified using the Qubit dsDNA HS Assay Kit (Thermo Fisher Scientific).

### Next generation library preparation

To evaluate the effects of variants, we employed the TileSeq approach (**Figure S3B**), in which the frequency of each variant is quantified in different sorted populations, such as BNP-high and BNP-low expression populations, via deep sequencing of small amplicons that tile the region of interest^49,64^. A three-step PCR protocol was performed for each tile to generate the sequencing library. All primers used in this study are provided in **Table S5**. Annealing temperatures and cycle times were optimized in advance.

PCR 1: Target Region Amplification

gDNA was extracted from the samples and amplified using primers binding outside the mutagenesis target region. All available gDNA was processed using 500 ng of DNA per 100 µL PCR reaction with KOD One PCR Master Mix (TOYOBO). Multiple PCR reactions were performed for each sample, pooled together, and purified using the QIAquick Gel Extraction Kit. Unedited iPSCs were processed in the same manner to serve as an unedited control.

PCR 2: Nested PCR with Illumina Adapters

A nested PCR was performed for each tile using 1 µL of purified PCR 1 product.

Primers containing Nextera adapters were designed for each tile to generate end-to-end coverage amplicons compatible with Illumina paired-end sequencing (2 × ∼150 bp tiles).

PCR 3: Dual Indexing and Library Preparation

Each PCR 2 product was dual-indexed by PCR using Nextera index primers. The resulting PCR products were purified using a PCR purification kit and quantified with the Qubit HS Assay Kit. Library preparation was carried out following Illumina guidelines, and sequencing was performed using an Illumina HiSeq platform.

Approximately 3 million reads were allocated per sample, and 30% PhiX control (Illumina) was included in the sequencing run.

The initial two CRISPR-X and ABE mutagenesis iPSC libraries were analyzed separately to confirm the introduction of variants and their distribution by each base editor (**Figure 2E**). Following amplification using 1st PCR primers, library preparation was performed according to the manufacturer’s protocol for the Nextera XT DNA Library Preparation Kit (Illumina) and Nextera XT Index Kit (Illumina). Approximately 1 million reads were allocated per sample, and sequencing was conducted using the Illumina iSeq platform with the inclusion of 20% PhiX control.

### Variant scoring

A custom pipeline was used to process sequencing data and generate variant counts. Variant counting at each position was performed using samtools mpileup after applying quality filtering for nucleotides with a base quality score over 30. The parental samples (e.g., BNP-high and BNP-low) were used to calculate the variant enrichment score at each base by comparing the percentage of reads with variants in the sample to the same percentage calculated in the parental sample. Sequencing data from the untreated control sample was used as a parental sample for the calculation of enrichment scores in the mutagenesis dataset.

1. The variant frequency in each replicate of the mutagenized iPSC library was less than 1x10^-4^.
2. The variant frequency was less than the estimated sequencing error calculated from the untreated control sample.
3. The variant was located in the primer binding region in each tile.
4. The variant was absent in both iPSC and iPSC-CM samples in the same replicate.
5. The variant frequency in each replicate of functional assay samples was less than 5x10^-4^.
6. The variant was not detected in at least two replicates in each functional assay.

We defined the Z-scored mean log2 enrichment score for each variant across replicates as the functional assay score for that variant. For variants located in overlapping regions of adjacent tiles, the mean of their scores was used as the final variant score. The score for each amino acid substitution was obtained by averaging the scores of variants that resulted in the same amino acid substitution.

### Classification of variants based on functional scores

First, we modeled the distributions of known pathogenic and benign variants using functional analysis data with kernel density estimation (KDE, **Figure S5E**, **S5G**, and **S5I**). The kernel density estimation was computed based on the following equation:

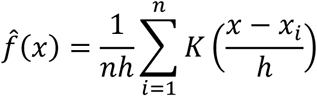

where represents the estimated density function, n is the number of data points, h is the bandwidth, and K is the kernel function. We utilized a Gaussian kernel, and the optimal bandwidth was determined using GridSearchCV with cross-validation, selecting the bandwidth that maximized the likelihood for each variant category (pathogenic and benign). To establish a classification cutoff, we calculated the log-likelihood ratio (LLR) as follows:

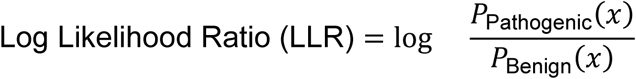

Where P_Pathoigenic_(x) and P_Bebign_(x) denote the probability density functions of the pathogenic and benign variant distributions, respectively. A positive LLR indicates a higher likelihood of pathogenicity, whereas a negative LLR suggests a higher likelihood of benignity.

The classification cutoff was defined as the zero-crossing point where the LLR transitions from negative to positive. If multiple zero-crossings were present, we selected the crossing point with the highest functional score, ensuring the boundary reflects the most pathogenic-associated region. This approach prevents choosing irrelevant intersections that do not effectively separate benign from pathogenic variants.

Furthermore, an intermediate classification zone was defined as the range of LLR within ± 0.5 around this cutoff, representing a region with lower classification confidence. Variants falling outside this intermediate range were classified as either pathogenic (higher scores) or benign (lower scores).

### OddsPath calculation

To estimate the strength of evidence for a given assay without performing rigorous statistical analysis, we calculated the odds of pathogenicity (OddsPath) for a theoretical assay by evaluating different numbers of known pathogenic and benign controls. The OddsPath calculations (OddsPath_Pathogenic and OddsPath_Benign) have been previously described^41^. We treated the proportion of pathogenic variants in the overall modeled data as the prior probability (P₁) and the proportion of pathogenic variants in the groups with functionally abnormal or functionally normal readouts as the posterior probability (P₂). The OddsPath was then calculated using the following formula:

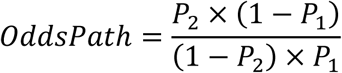

Each calculated OddsPath was then applied to a corresponding level of evidence strength according to the Bayesian adaptation of the ACMG/AMP variant interpretation guidelines.

### Protein 3D structure and structural analysis

Protein structures from the Protein Data Bank (PDB) were visualized using PyMOL (v3.1.3). The NMR structure (PDB: 5K6P)^65^ and the electron microscopy structure (PDB: 7TIJ)^36^ of the cMyBP-C M-domain were used. For the 3D structure shown in **Figure 5E**, the corresponding region of the AlphaFold-predicted structure was used to represent THB and the disordered region. In protein stability analysis, images of the protein structure were made using YASARA View (v25.1.13)^66^ on the structure of MYBPC3 from PDB 5K6P, chain A, object 1^65^. The **ΔΔ**G calculations were made using FoldX (v5.1)^38,39^. The median **ΔΔ**G was reported across each of the 20 NMR objects in 5K6P (**Figure S7**).

## SUPPLEMENTAL FIGURES

**Figure S1.**
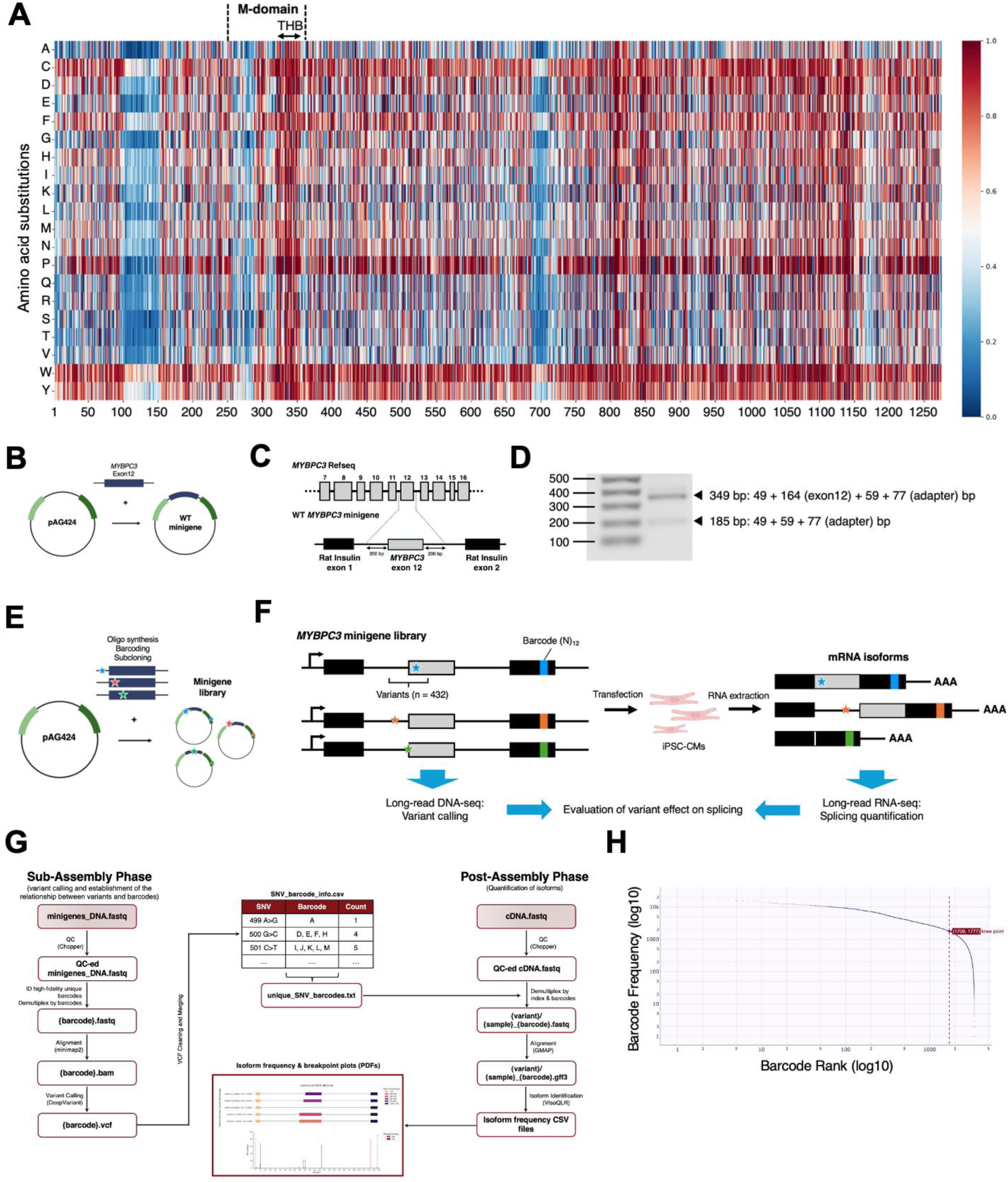
AlphaMissense prediction of cMyBP-C and preparation for minigene splicing library. **A,** AlphaMissense prediction scores in cMyBP-C. **B,** Schematic of reference sequence minigene design and cloning. Created with BioRender.com. **C,** *MYBPC3* reference sequence minigene consisting of *MYBPC3* exon 12 (164bp) and 200 bp of native intronic flanking regions on both sides of the exon (intron 11 and 12), flanked by rat insulin exons 1 and 2. **D,** Reference sequence minigene generated the same splicing isoform (the canonical isoform) as endogenous *MYBPC3.* **E,** Design of minigene library. Oligo synthesis, barcoding, and cloning were performed by Twist Bioscience. Created with BioRender.com. **F,** Massively parallel splicing assay using minigene library. The minigene library includes 432 variants at the intron 11-exon 12 junction. Each plasmid was tagged with a 12-nt unique barcode. Variants and splicing isoforms are characterized by long DNA and RNA sequencing and linked by unique barcode information. Created with BioRender.com. **G,** The workflow of custom pipeline to evaluate the splicing assay results. **H,** Knee plot for the barcodes quality filtering in minigene library. 5201 unique barcodes were identified in DNA_fastq after filtering.

**Figure S2.**
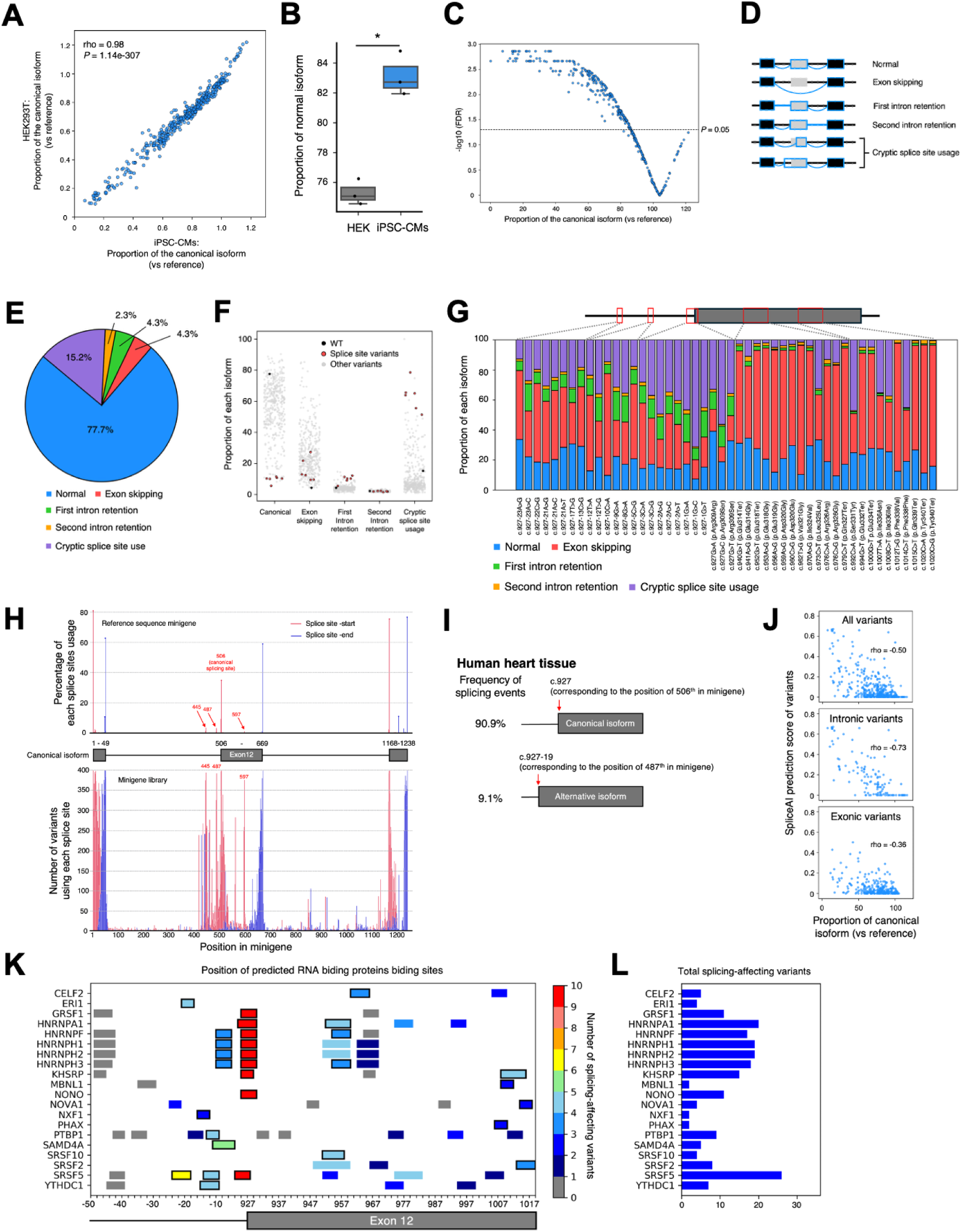
Minigene splicing assay to assess variant effects on splicing. **A,** Correlation of normalized frequency of canonical isoform in HEK293T and iPSC-CMs. **B,** Proportion of the canonical isoform of reference minigene in HEK293T cells (n = 3) and iPSC-CMs (n = 3). The proportion of the canonical isoform in iPSC-CMs was slightly higher than in HEK293T cells (HEK293T; 75.2 ± 0.5%, iPSC-CMs; 83.1 ± 0.9%, *p* = 1.35e-3). **C,** Volcano plot of normalized frequency of canonical isoform and -log10 (FDR). FDR indicates false discovery rate. There were 14 splice-neutral variants located in the intron at a location enriched for predicted splice factor binding that increased the proportion of the canonical isoform observed compared to reference sequence, although these did not reach statistical significance. **D,** Classification of splicing isoforms. 115 distinct splicing isoforms were identified in the minigene library. Isoforms were classified as canonical, exon skipping, first intron retention, second intron retention, or cryptic splice site usage. **E,** Proportion of each isoform in reference sequence minigene. **F,** Proportion of each isoform class across 402 variants. **G,** Splice site usage in reference sequence minigene (top) and minigene library (bottom). Several common alternative splicing sites were identified in reference and other variants. **H,** Alternative splice site identified in minigene assay was also observed in human heart tissue. **I,** Splicing outcome of splice-affecting variants (variants classified as intermediate and disrupting). The isoform frequencies are shown as mean values from 4 biological replicates. Cryptic splice site usage was primarily driven by intronic variants, especially those located at canonical splice sites, while exon skipping was observed as a splicing outcome for both intronic and exonic variants. **J,** Correlation between splicing assay scores and SpliceAI prediction scores (All variants; rho = -0.50, *p* = 4.5e-27, Intronic variants; rho = -0.72, *p* = 5.2e-21, Exonic variants; rho = -0.36, *p* = 7.6e-10). The SpliceAI scores range from 0 to 1, where higher scores indicate a greater probability that a variant alters splicing. **K,** The 20 RNA binding proteins (RBPs) are predicted to bind in *MYBPC3* intron 11 and exon 12 junction. The active RBP binding sites (more than 50% of positions were associated with splicing-affecting variants) are highlighted. **L,** Total splicing-affecting variants located at each RBPs binding sites.

**Figure S3.**
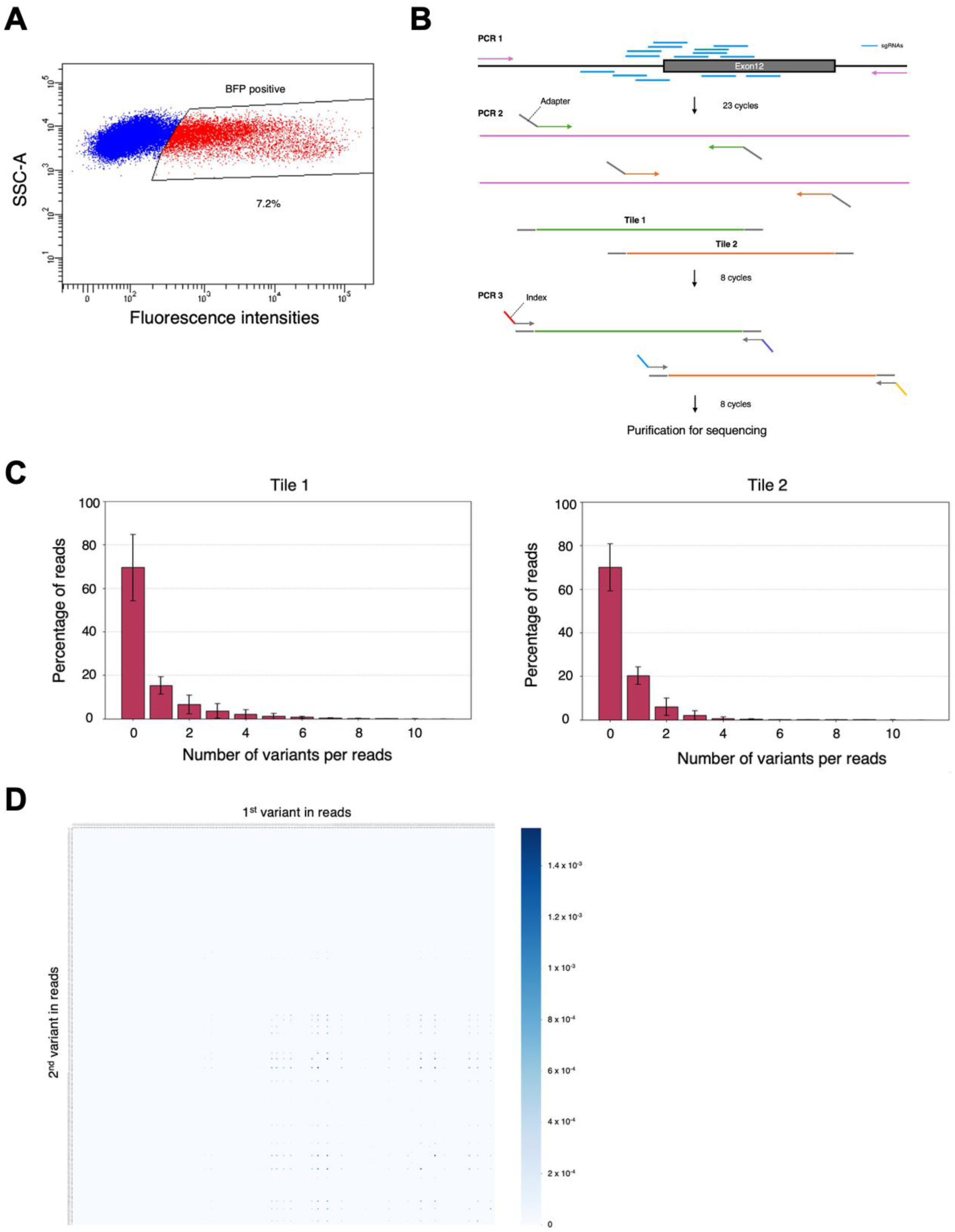
*in situ* mutagenesis and bystander editing. **A,** Following electroporation of base editor and pooled sgRNA, BFP+ iPSCs were sorted by FACS to enrich mutagenized iPSCs. **B,** Frequency of variants were quantified via amplicon sequencing. **C,** Distribution of variant counts per read. **D,** Variant pairs frequency in reads.

**Figure S4.**
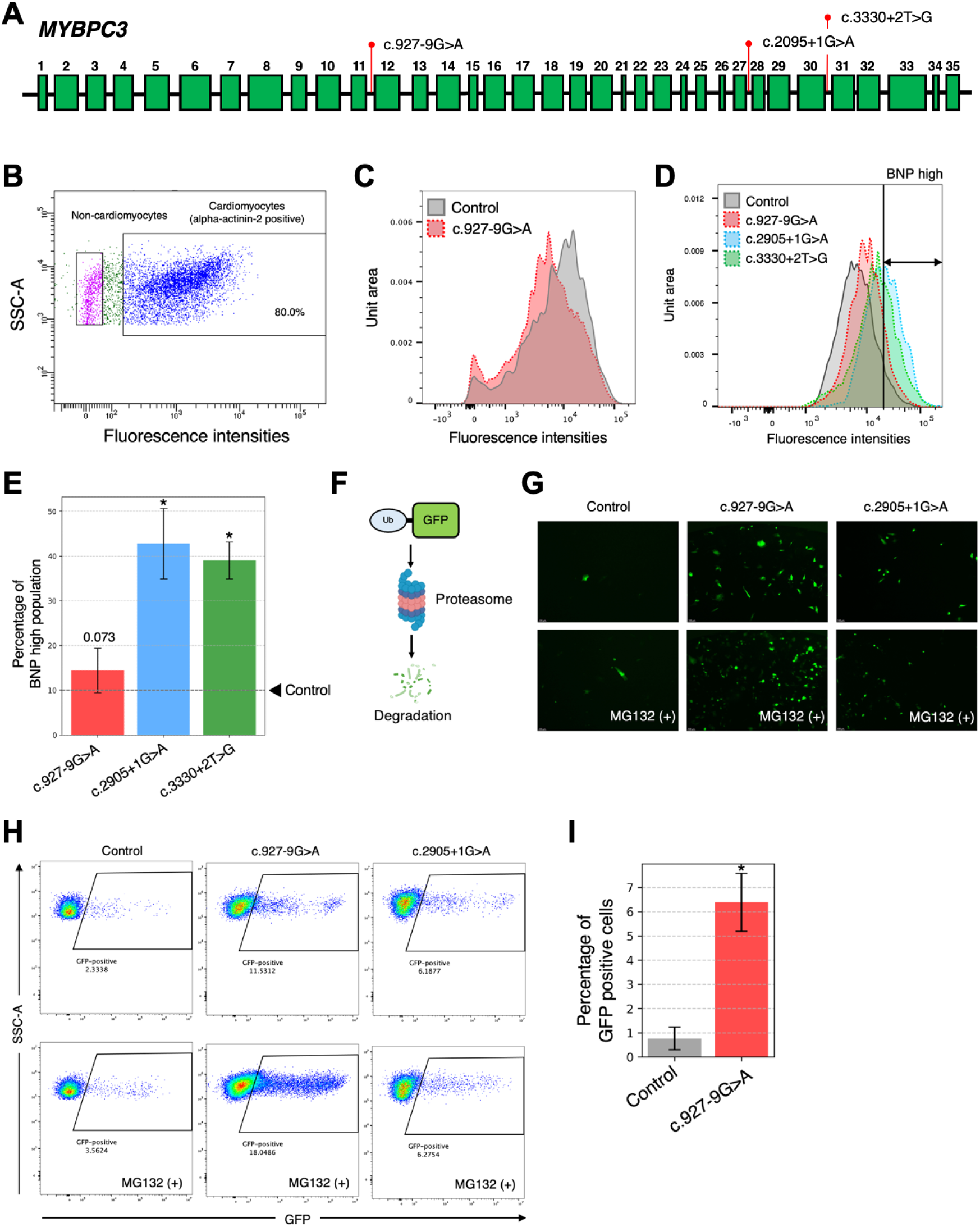
Functional assays using pathogenic *MYBPC3* iPSC-CMs. **A,** Pathogenic variants used for preliminary functional assays. **B,** iPSC-CMs were sorted based on alpha-actinin-2 expression by fluorescence-activated cell sorting (FACS). **C,** Distribution of cMyBP-C abundance level in control iPSC-CMs and iPSC-CMs carrying c.927-9G>A. **D,** Distribution of BNP expression level in control iPSC-CMs and iPSC-CMs carrying pathogenic *MYBPC3* variants. **E,** iPSC-CMs carrying c.2905+1G>A and c.3330+2T>G exhibited significantly higher BNP expression compared to control iPSC-CMs. (n = 3-4, \**p* < 0.05) **F,** Ubiquitin fusion degradation targeted GFP (Ub-GFP) allows quantification of ubiquitin proteasome system activity in living cells. Created with BioRender.com. **G** and **H,** GFP expression in iPSC-CMs 48 hrs after transfection of Ub-GFP. MG132 is a proteasome inhibitor (50μM). **I,** iPSC-CMs carrying c.927-9G>A showed a significantly higher percentage of GFP positive cells compared to control (n = 3).

**Figure S5.**
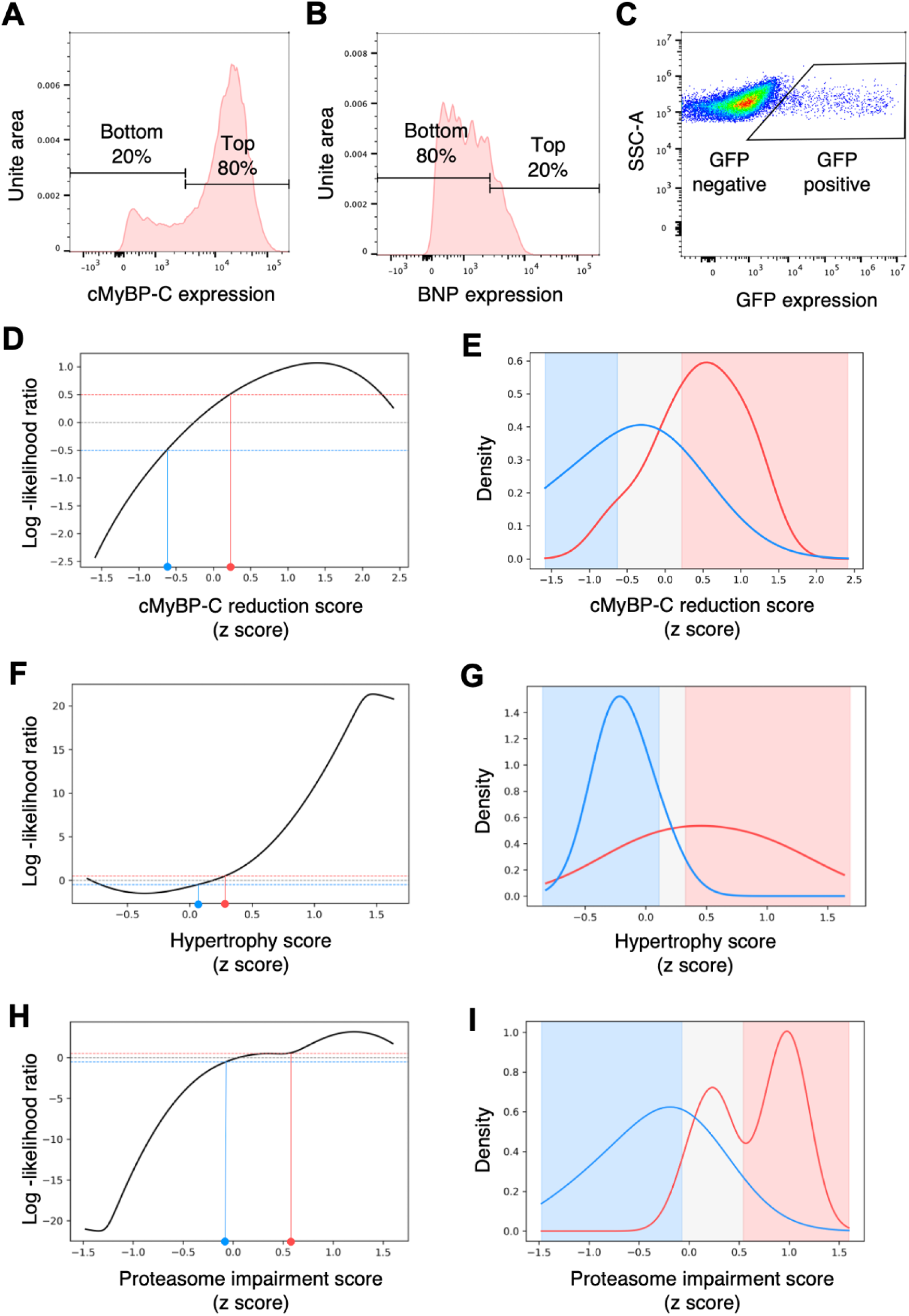
Functional assays using mutagenized iPSC-CMs library. **A** through **C,** Sorting strategy to evaluate variant effects in each assay. **D** through **I,** To classify variants based on their functional scores, log-likelihood ratios (LLRs) were calculated from well-established pathogenic or benign variants and used to set thresholds. An LLR range from -0.5 to 0.5 was defined as the intermediate range (light gray). The benign range was indicated in light blue, and the pathogenic range in pink.

**Figure S6.**
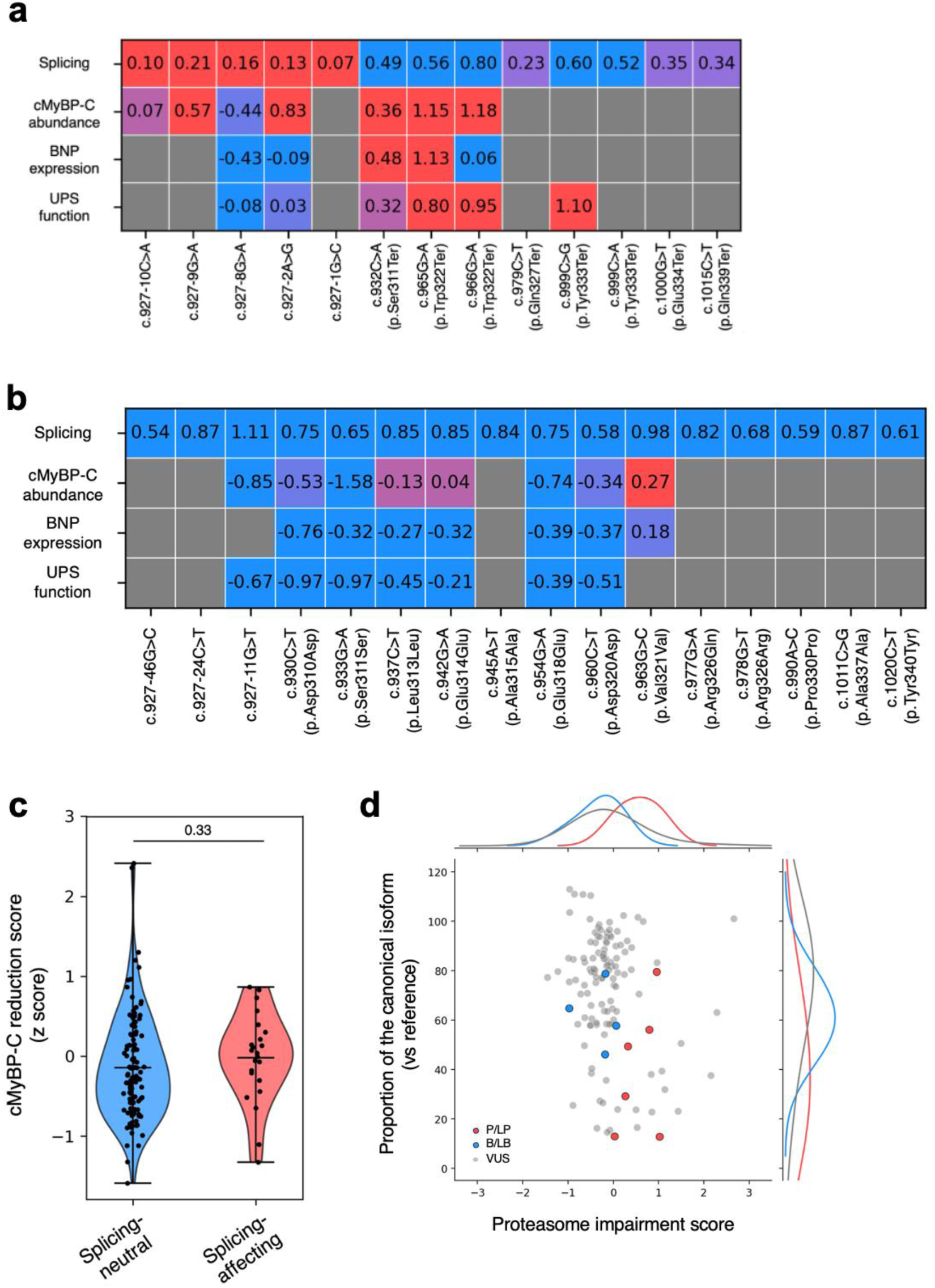
Integrated analysis of multidimensional functional assay. **A,** Functional scores in pathogenic or likely pathogenic variants in ClinVar. **B,** Functional scores in benign or likely benign variants in ClinVar. **C,** Protein reduction score in splicing-neutral and -affecting variants. **D,** Comparison of normalized proportion of the canonical splicing isoform and proteasome impairment scores.

**Figure S7.**
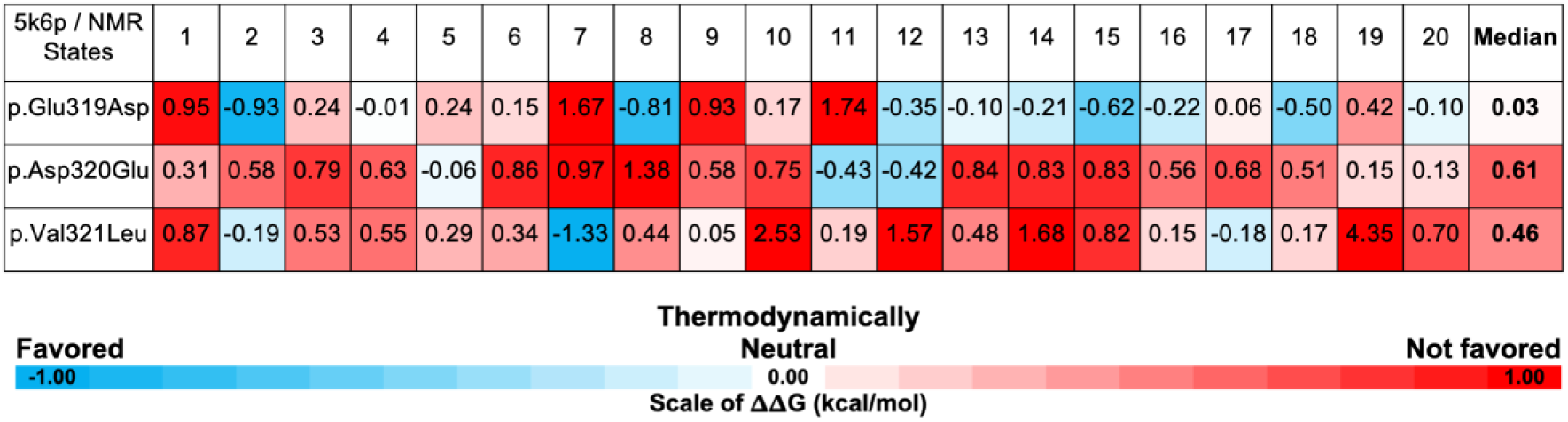
Assessment of protein stability through f ΔΔG (kcal/mol).

## SUPPLEMENTAL TABLES

**Table S2.** Known pathogenic/likely pathogenic and benign/likely benign variants.

Pathogenic
| Genomic allele | cDNA variant | Protein variant | Variant consequence |
| --- | --- | --- | --- |
| 11-47346379:C:T | c.927-9G>A |  | INTRONIC |
| 11-47346372:T:G | c.927-2A>C |  | CANONICAL_SPLICE |
| 11-47346372:T:C | c.927-2A>G |  | CANONICAL_SPLICE |
| 11-47346372:T:A | c.927-2A>T |  | CANONICAL_SPLICE |
| 11-47346371:C:T | c.927-1G>A |  | CANONICAL_SPLICE |
| 11-47346371:C:G | c.927-1G>C |  | CANONICAL_SPLICE |
| 11-47346371:C:A | c.927-1G>T |  | CANONICAL_SPLICE |
| 11-47346365:G:T | c.932C>A | p.Ser311Ter | STOP_GAINED |
| 11-47346363:T:A | c.934A>T | p.Lys312Ter | STOP_GAINED |
| 11-47346357:C:A | c.940G>T | p.Glu314Ter | STOP_GAINED |
| 11-47346345:C:A | c.952G>T | p.Glu318Ter | STOP_GAINED |
| 11-47346342:C:A | c.955G>T | p.Glu319Ter | STOP_GAINED |
| 11-47346332:C:T | c.965G>A | p.Trp322Ter | STOP_GAINED |
| 11-47346331:C:T | c.966G>A | p.Trp322Ter | STOP_GAINED |
| 11-47346330:C:A | c.967G>T | p.Glu323Ter | STOP_GAINED |
| 11-47346318:G:A | c.979C>T | p.Gln327Ter | STOP_GAINED |
| 11-47346303:C:A | c.994G>T | p.Glu332Ter | STOP_GAINED |
| 11-47346298:G:T | c.999C>A | p.Tyr333Ter | STOP_GAINED |
| 11-47346298:G:C | c.999C>G | p.Tyr333Ter | STOP_GAINED |
| 11-47346297:C:A | c.1000G>T | p.Glu334Ter | STOP_GAINED |
| 11-47346282:G:A | c.1015C>T | p.Gln339Ter | STOP_GAINED |
| 11-47346277:G:T | c.1020C>A | p.Tyr340Ter | STOP_GAINED |
| 11-47346277:G:C | c.1020C>G | p.Tyr340Ter | STOP_GAINED |

Benign
| Genomic allele | cDNA variant | Protein variant | Variant consequence |
| --- | --- | --- | --- |
| 11-47346416:C:G | c.927-46G>C |  | INTRONIC |
| 11-47346398:G:A | c.927-28C>T |  | INTRONIC |
| 11-47346397:C:G | c.927-27G>C |  | INTRONIC |
| 11-47346364:C:T | c.933G>A | p.Ser311Ser | SYNONYMOUS |
| 11-47346364:C:G | c.933G>C | p.Ser311Ser | SYNONYMOUS |
| 11-47346336:C:T | c.961G>A | p.Val321Met | NON_SYNONYMOUS |
| 11-47346320:C:T | c.977G>A | p.Arg326Gln | NON_SYNONYMOUS |
| 11-47346297:C:T | c.1000G>A | p.Glu334Lys | NON_SYNONYMOUS |
| 11-47346277:G:A | c.1020C>T | p.Tyr340Tyr | SYNONYMOUS |

**Table S3.**
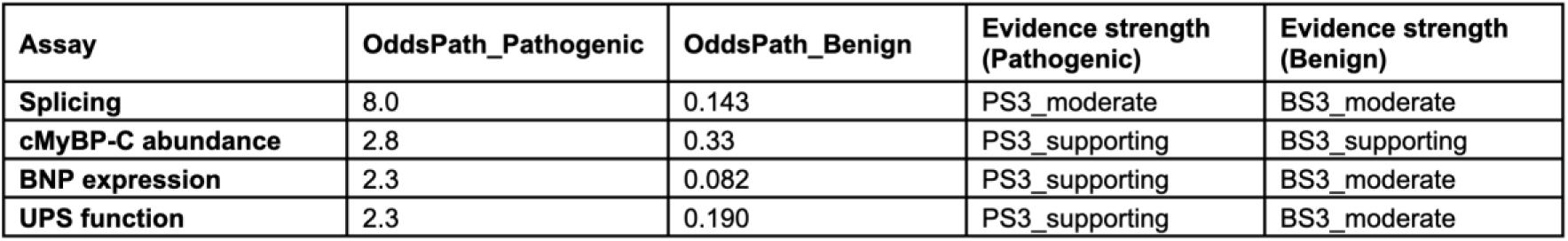
Odds of pathogenicity.

**Table S4.**
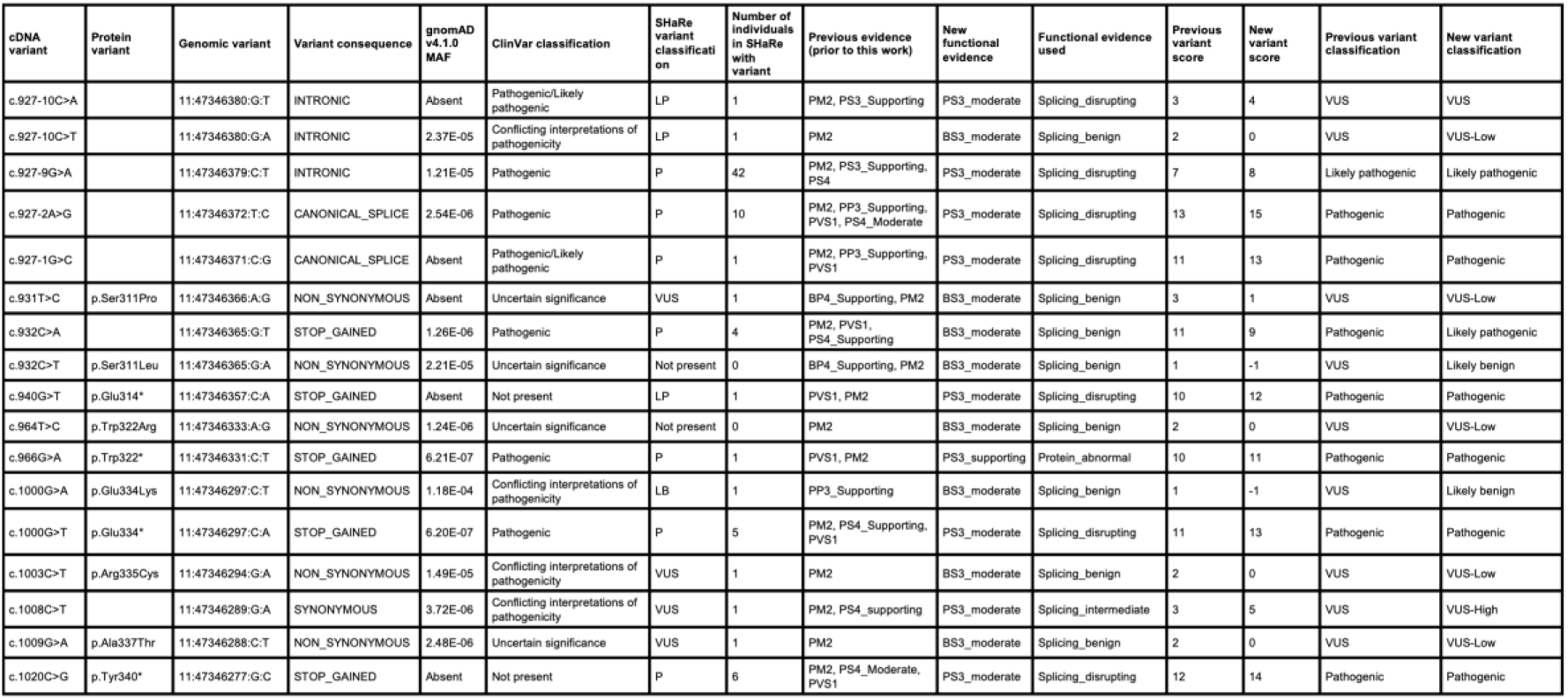
*MYBPC3* variants in SHaRe registry.

